# Oncogenic mutations disrupt lineage differentiation via epigenetic perturbations

**DOI:** 10.1101/2025.04.22.649974

**Authors:** Lingjun Meng, Alice Yu, Huanwei Huang, Jodie Meng, Yongjin Yoo, Katie Schaukowitch, Justyna A. Janas, David A Knowles

## Abstract

Understanding lineage differentiation is an ongoing challenge in the field of stem cell research. We acquired eight gain-of-function (GOF) oncogenic mutations associated with DNA methylation alteration from the TCGA datasets. After introducing each oncogene into wild-type stem cells separately, we found that the mutant stem cells are difficult to reprogram to iPSCs and terminally differentiate to induced neuronal cells, and retain proliferation capacity in the process of differentiation. This findings suggests that oncogenic mutations impair the lineage differentiation. After analyzing the resulting DNA methylation profiles, we identified many novel lineage differentiation genes in mutant cells showing DNA methylation alteration compared with wild-type cells, which is consistent with their methylation states in low-grade gliomas (LGGs) compared with control brain. Collectively, our results indicate that oncogenic mutations in stem cells are associated with DNA methylation changes for genes related to multipotency maintenance, differentiation resistance, and tumorigenesis, therefore, displaying impairment in lineage commitment.

## Introduction

Induced pluripotent stem cell (iPSC) reprogramming and cell differentiation hold great promise for regenerative medicine. However, their potential clinical application is hampered by the low efficiency of somatic cell reprogramming and differentiation. We previously found that cancer cells are extremely difficult to reprogram into iPSCs, except in acute myeloid leukemia (AML)^1^. Reprogramming resets epigenetic states in AML-iPSCs, but leukemic behavior and methylation are reacquired upon hematopoietic differentiation^1^. There appears to be a specific epigenetic profile related to cell behavior during lineage differentiation. A better understanding of oncogenic mutation associated epigenetic profiles holds great promise for enhancing reprogramming and differentiation efficiency. Here, we show that oncogenic mutations interrupt iPSC reprogramming and neural cell differentiation via change of epigenetic states, especially reveals that methylation changes appear on genes play a critical role on multipotency, differentiation, and tumorigenesis.

By analyzing DNA methylation profiles across cancers represented in TCGA (The Cancer Genome Atlas), we identified eight candidate oncogenic mutations that are most associated with genome-wide DNA methylation changes. To investigate the effect of oncogenic mutations in reprogramming and differentiation, we independently introduced each oncogene into wild-type (wt) neural progenitor cells (NPCs) derived from human embryonic stem cells (hESCs) and investigated the epigenetic alteration occurring after the induction of the oncogenes. We found that the induction of gain-of-function mutation of oncogenes effects the change of the epigenetic landscape in mutant NPCs compared to wt NPCs. To understand the role of oncogenes in lineage commitment, we measured the differentiation efficiency of hESCs (or NPCs) to induced neuronal cells (iN), and the reprogramming efficiency of mutant NPCs to human induced pluripotent stem cells (hiPSCs). Through comparing the efficiency of mutant cells with wt cells in the process of lineage reprogramming and differentiation, we found the mutant cells displayed a preference for remaining as stem cells and resisting lineage change. Moreover, the mutant cells retained their proliferation capacity in dysregulated lineage commitment and consequently generated more cells in a differentiation resistance status due to inaccurate lineage commitment. Through comparing the DNA methylation change between wt cells and mutant cells before and after lineage differentiation, we found that many genes that play a role in multipotency maintenance and neural identification had altered methylation, consistent with the same methylation states in low-grade gliomas (LGGs). Our results indicate that the significant role of oncogenes which is associated with the expression of genes that have a function on lineage commitment through epigenetic modification.

## Results

### Expression of oncogenic mutations blocks differentiation of ES cell into neurons

It was previously shown that IDH1 (isocitrate dehydrogenase 1) mutations occur in the majority of gliomas^2^. IDH1 mutant cells generate R(2)-2-hydroxyglutarate (2HG)^3^, which causes GASC1 (lysine-specific demethylase 4C, KDM4C) mediated hypermethylation^4,5^. Building upon these previous studies^2,4^, we chose the IDH1-mutation as our first candidate to study the correlation between oncogene and epigenetic alteration. We investigatd the frequency of IDH1 mutation within each cancer type in TCGA. We confirmed that IDH1 mutation frequently occurs in low-grade gliomas (LGGs) and glioblastoma (GBM) (Extended Data Fig. 1a). Thus, we interrogated the effect of IDH1-R132* mutations (R132C, R132G, R132H, R132S) on the epigenetic states in GBM and LGG, as measured by DNA methylation. IDH1-R132 mutations are highly associated with DNA methylation changes (Extended Data Fig. 1b, c).

To understand whether oncogenes other than IDH1 show similar patterns, we performed a pan-cancer analysis linking both gain-of-function (GoF) and loss-of-function (LoF) mutation changes to epigenetic state, as represented by principal components of DNA methylation. Across The Cancer Genome Atlas, we detected 22 and 17 candidate genes associations of DNA methylation for GoF and LoF mutations respectively (1% false discovery rate) (Extended Data Fig. 1d, e). We picked the top eight gain-of-function genes (IDH1-R132H, TP53-R273C, BRAF-V600E, KRAS-G12C, NRAS-Q61R, HRAS-Q61R, SOX17-S403I, FGFR3-S294C) from the candidate list based on p-value (adjusted pval < 0.05) for the following experimental validation. It indicated that not only the IDH1 mutation reported before, but also a range of oncogenic mutations are associated with epigenetic alteration in pan-cancer DNA methylation analysis.

To investigate the function of the eight oncogenes indicated above in cell lineage commitment, we assessed the differentiation efficiency from mutant human embryonic stem cells (hESCs) to induced neuronal (iN) cells. We used a doxycycline (dox)-inducible lentiviral vector system to transduce empty vector plasmid as a control and eight candidate oncogenes into hESC H1 cells, separately. As we showed previously, overexpression of neuronal gene neurogenin-2 (Ngn2) efficiently converted hESCs into iN cells with nearly 100% yield and purity in less than two weeks ^6^ (Fig. 1a). We compared the Ngn2 induced differentiation efficiency between wt and mutant iN cells by staining with neuronal marker Tuj1 (tubulin beta-III). The differentiation efficiency of wt hESCs is in a significant high percentage as nearly 100%. The wt hESC-derived iN cells showed typical neuronal morphology with extension of axons and smaller cell nuclei. The differentiation of iN cells from hESCs forces the cells to completely lose their stem cell identification and proliferation capacity. Remarkably, for all eight tested candidate oncogenes, a high percentage of mutant hESC cells failed to differentiate to iN cells (Fig. 1b). We demonstrated that mutant hESC-derived iN cells have immature iN cell body morphology and short axons (Fig. 1b). Their nuclear size is similar to hESCs, and not as small as wt hESC-derived iN cells (Fig. 1b). We measured neurite length of cells at 12 days of iN differentiation and found markedly shorter neurites in mutant hESC-derived iN cells than in wt hESC-derived iN cells (Fig. 1c). In addition, we see a reduction in mRNA expression level of the neuronal marker Map2 in mutant hESC-derived iN cells compared to wt hESC-derived iN cells by qRT-PCR analysis at day 7 which is the early stage of differentiation (Fig. 1d).

**Fig. 1.**
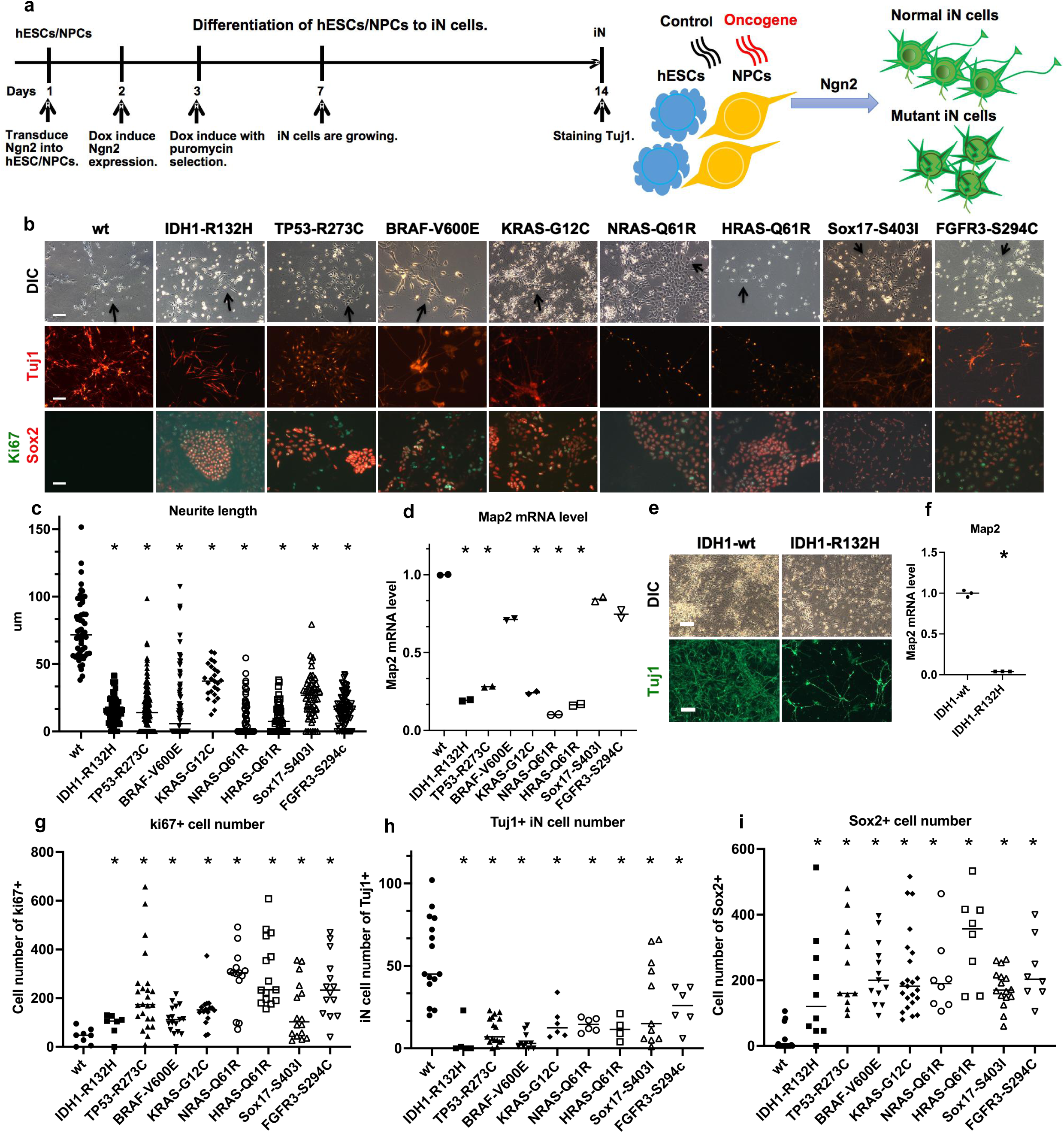
Expression of oncogenic mutations blocks differentiation of ES cell into neurons. **a,** Experimental scheme for generating induced neuronal (iN) cells differentiated from human embryonic stem cells (hESCs) or neural progenitor cells (NPCs) by overexpression of the neuronal gene, Ngn2. **b,** Brightfield and fluorescent images of iN cells stained for the neural cell marker Tuj1 (red), pluripotency marker Sox2 (red), cell proliferation marker Ki67 (green) from wt hESC-derived iN cells and mutant hESC-derived iN cells after 2 weeks of differentiation. Data are from at least three independent experiments. Scale bars, 50 μm. **c,** Neurite length in cells were measured at 12 days of iN differentiation (*p < 0.01). **d,** Quantitative RT-PCR analysis of neural cell marker Map2 expression level at 7 days of hESC differentiation. Two-tailed t-test, Mean ± SD (n=3, *p < 0.01). **e,** Brightfield and fluorescent images of NPC-derived iN cells stained for the neural cells marker Tuj1 (green) from IDH1-wt NPC-derived iN cells and IDH1-R132H mutation NPC-derived iN cells after 2 weeks of differentiation. **f,** Quantitative RT-PCR analysis of neural cell marker Map2 expression at 7 days of NPC differentiation. Values represent mean ± SD (n=3, *p < 0.01). **g,** Cell number counting of Ki67 positive wt hESC-derived iN cells and mutant hESC-derived iN cells at 10 days of differentiation (*p < 0.01). **h,** Cell number counting of Tuj1 positive wt hESC-derived iN cells and mutant hESC-derived iN cells at 10 days of differentiation (*p < 0.01). **i,** Cell number counting of Sox2 positive wt hESC-derived iN cells and mutant hESC-derived iN cells at 10 days of differentiation. Values represent the number of cells showing positive in each situation ± SD (n=3, *p < 0.01).

Consistent with IDH1-R132H mutant hESCs, the IDH1-R132H mutant NPCs are also hard to differentiate to iN cells compared to IDH1-wt NPCs (overexpress IDH1 without any mutations) (Fig. 1e). From qRT-PCR analysis of neural marker expression at day 7 of differentiation from NPCs, we found a significant decrease in the expression level of Map2, Nestin, Pax6, and NeuroD1 in IDH1-R132H mutation NPC-derived iN cells compared to IDH1-wt NPC-derived iN cells (Fig. 1f, Extended Data Fig. 2a, b, c). Collectively, these results suggest that oncogenic mutations cause difficulty in inducing terminal differentiation from hESCs and NPCs to neuronal cells, and perturb cell fate determination.

As known, the terminally differentiated cells, such as the neuronal cells discussed here, cannot proliferate. Interestingly, we noticed that the mutant stem cells not only retain their stem cell state via maintaining the expression of Sox2 and resist neural differentiation, but they also continue to proliferate throughout the differentiation process. To investigate further, we counted the cell number of wt and mutant hESCs with dox induction for 3 days, but did not find an increase in mutant hESCs compared to wt hESCs (Extended Data Fig. 2d). This suggested that there is no proliferation advantage of mutant hESCs. However, we found that the cell number of mutant hESC-derived iN cells was increased after dox for 7 days compared to wt hESC-derived iN cells (Extended Data Fig. 2e). To validate the possibility that oncogenes may enhance cell proliferation, we stained the wt and mutant hESC-derived iN cells with the cell proliferation marker Ki67 (Fig. 1g). For all eight oncogenes, we found that a high percentage of mutant hESC-derived iN cells express Ki67 during the differentiation process, but rarely happen in wt hESC-derived iN cells. To identify the cell type of Ki67+ cells, we co-stained for the neuronal marker Tuj1 and pluripotency marker Sox2 with Ki67. There were three types of cells: Ki67+Sox2+, Ki67+Sox2– and Ki67+Tuj1–. Ki67+Tuj1+ cells were never observed. We counted the Ki67+ cells at 10 days of differentiation. Very few wt hESC-derived iN cells were Ki67+, but more than hundreds times mutant hESC-derived iN cells retained their proliferation capacity and resisted being fully differentiated to neural cells (Fig. 1g). There was a significant reduction in neuronal cell number of mutant hESC-derived iN cells at day 10 (Fig. 1h), even when the mutant hESC-derived iN cells expressed Tuj1, most of them did not grow long axons. There were hundreds times more Sox2+ cells in the eight mutant hESC-derived iN cells than wt hESC-derived iN cells at 10 days of differentiation (Fig. 1i, Extended Data Fig. 2f). In summary, we found that stem cells expressing oncogenes enable themselves to resist differentiation into neural cells. This suggests that many oncogenic mutations may be associated with inhibition of differentiation potential due to epigenetic rewiring, because it has only happened during lineage differentiation with more permissive epigenetic states.

### Stem cells with oncogenic mutations are difficult to reprogram into iPS cells

As it was previously found that cancer cells are extremely difficult to reprogram into iPSCs^1^, we employed human neural progenitor cells (NPCs) as above used, and planned to assess the human iPSC (hiPSC) reprogramming efficiency from NPCs with these eight oncogenes shown in Fig 1, in order to comprehensively understand the lineage commitment blocking. We used NPCs derived from hESCs following the dual smad inhibition protocol ^7^ (Extended Data Fig. 3). To investigate the effect of each oncogene, we used the dox inducible system, same as discussed above. The NPCs were used to generate hiPSCs by overexpressing the four transcription factors: Oct4, Sox2, c-Myc, and Klf4 (OSKM)^8^ (Fig. 2a). Afterward, the reprogramming efficiency of wt and mutant NPC-derived iPSCs were compared by counting the number of iPSC colonies via immunofluorescence staining of an iPSC marker, TRA-1-60 (Fig. 2b, c). Wt and IDH1-wt NPC-derived iPSCs generated many colonies. In contrast, all eight mutant NPCs were difficult to reprogram into iPSCs and generated no iPSC colonies or a few very small colonies (Fig. 2b, c). These results were consistent with our previous study showing that cancer cells were difficult to reprogram to iPSCs (Extended Data Fig. 4), even if there was only one oncogene induced in stem cells with a short-term inducible expression. In summary, stem cells with oncogenes are difficult to reprogram into iPS cells.

**Fig. 2.**
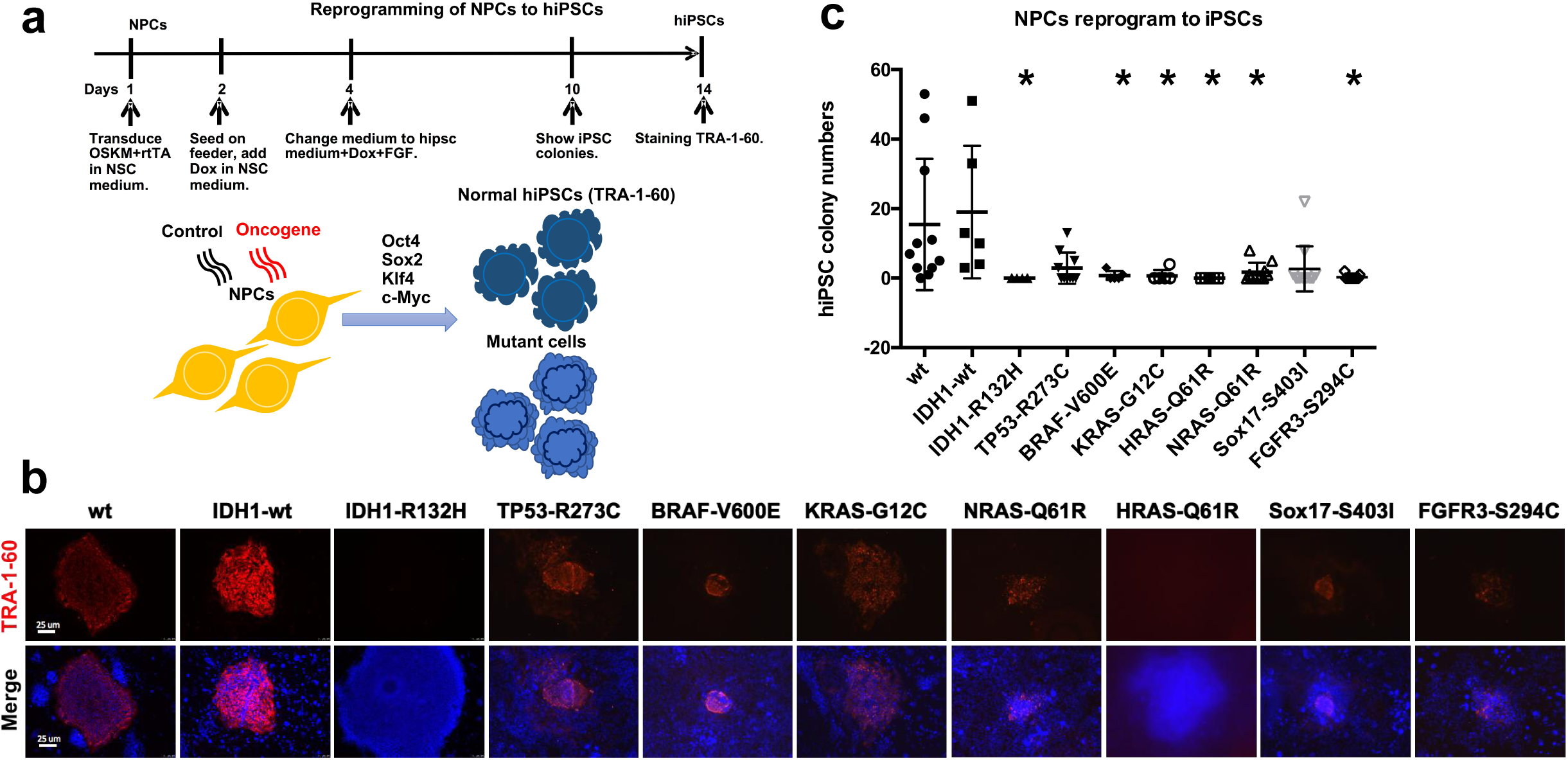
Stem cells with oncogenic mutations are difficult to reprogram into iPS cells. **a,** Experimental scheme for generating human induced pluripotent stem cells (hiPSCs) from human NPCs by overexpression of Oct4, Sox2, Klf4, and c-Myc (OSKM) four transcription factors. **b,** The fluorescent images of NPC-derived iPSCs stained for the pluripotency marker TRA-1-60 (red) from wt NPC-derived iPSCs and mutant NPC-derived iPSCs. Nuclear staining by DAPI (blue). Scale bars, 25 μm. **c,** The number of TRA-1-60 positive iPSCs colonies generated from the reprogramming of wt and mutant NPCs (n=3, *p < 0.05).

### Oncogenic mutation is associated with epigenetic change in hESCs

To investigate the effect of oncogenic mutations on genome-wide DNA methylation in stem cells, we performed an Infinium MethylationEPIC BeadChip assays. To clarify that the epigenetic change is caused only by one oncogenic mutation, we introduced each oncogene into individual wt hESCs, the same as we previously did, using dox to overexpress the GoF mutation gene for nine days. The principal component analysis (PCA) of DNA methylation profiles demonstrated that the seven mutant hESCs and wt hESCs as control (empty vector) were distinguished by DNA methylated gene loci (Extended Data Fig. 5a, b). 282 CpG sites are differentially methylated in the seven mutants compared with control, 246 out of 286 are associated with specific genes (Extended Data Fig. 5c). TP53I3 and PTPRT were the top two genes differentially methylated in differentially methylated positions (DMPs) distribution. Furthermore, 321 genomic regions exhibited significant differences in differentially methylated regions (DMRs) distribution (p-val < 0.05) (Extended Data Fig. 5d). Based on the distance from CpG islands, there are 401 genomic regions that are differentially methylated in total 4,905 regions (p-val < 0.05) (Extended Data Fig. 5e). Geneset enrichment analysis showed that many tumor suppressor genes are methylated within DMPs and DMRs (Extended Data Fig. 5f, g). The genes are enriched for stem-like genesets, such as Wnt signaling pathway (WNT5A, UBC, WWTR1, Sox9), embryonic skeletal system morphogenesis (HOXB1, HOXA7), beta-catenin-TCF complex assembly (HIST1H4B, HIST1H4C), and positive regulation of transcription from RNA polymerase II promoter (ZBTB38). These findings demonstrated that oncogenic mutations are associated with epigenetic alteration in hESCs.

### Oncogenic mutations block differentiation via epigenetic alteration of stem cell fate genes

As shown above, stem cells with oncogenes were difficult to induce into terminal differentiation from NPCs to iN cells, which may be associated with epigenetic changes (Fig.1). To investigate temporal DNA methylation changes occurring during NPC maintenance in early and later times via genomic DNA methylation measurement by Infinium MethylationEPIC BeadChip assay, we introduced IDH1-R132H, KRAS-G12C, TP53-R273C, BRAF-V600E, NRAS-Q61R, HRAS-Q61R, and SOX17-S403I mutations separately into NPCs at day 0, with dox induction for 2 or 10 days to overexpress the oncogenes. After 10 days of induction, we differentiated the NPCs into iN cells for two more days using the same procedure described in Fig 1 (Fig. 3a), in order to investigate the epigenetic change during differentiation at increase temporal resolution. As a control, we conducted the same analysis on wt-2 days, wt-10 days, and wt-10 days + 2 iN days (Fig. 3a).

**Fig. 3.**
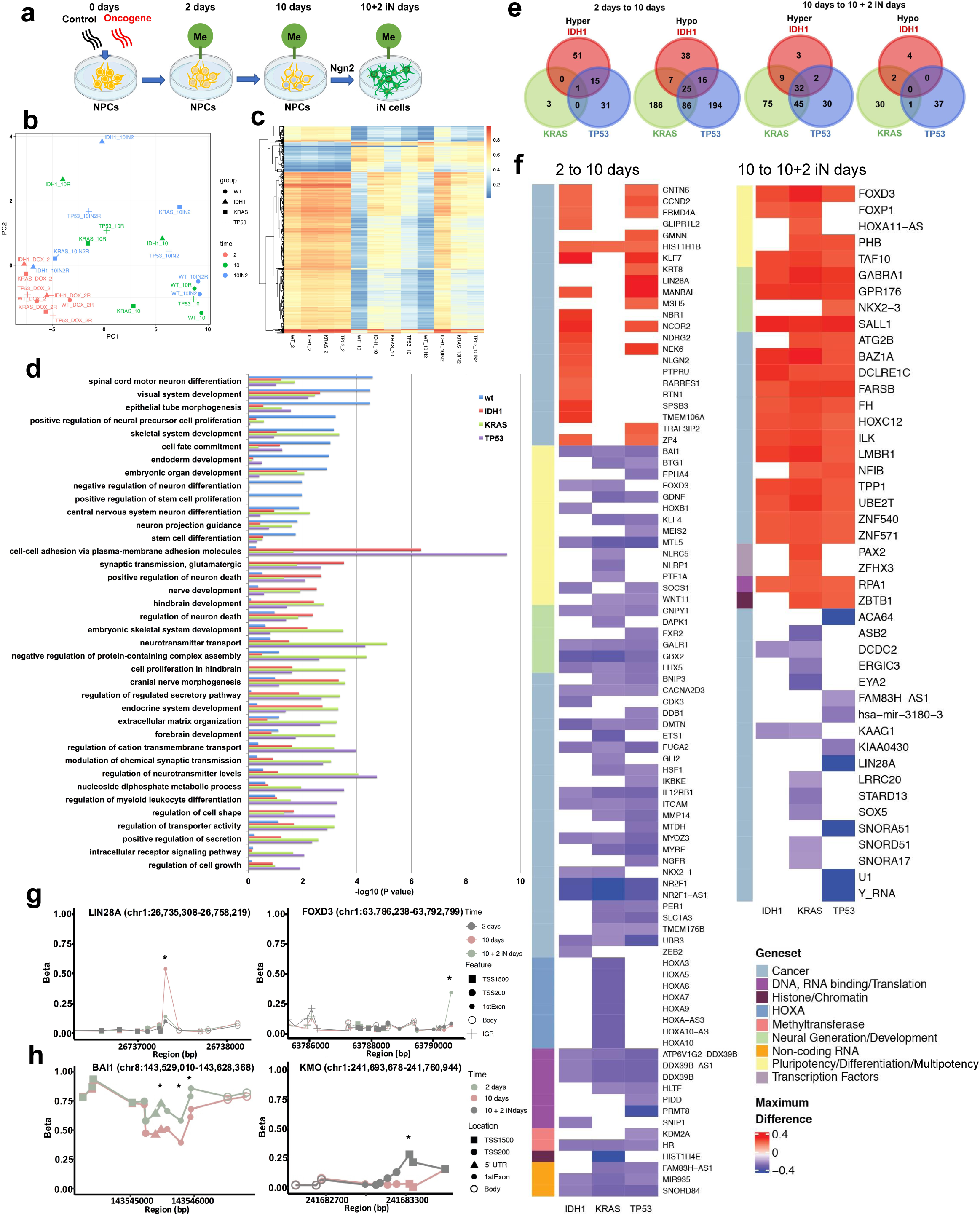
Oncogenic mutations block differentiation via epigenetic alteration of stem cell fate genes. **a,** A schematic timeline representation detailing experimental design. **b,** Principal component analysis (PCA) of DNA methylation profiles of seven oncogenes and wt NPCs are distinguished by DNA methylation states, from induced genes express 2 days, 10 days and 10 days plus 2 days induce to iN cells. PCA plot of the different methylation samples color-coded in time of induction (n=2). **c,** Heatmap analysis of significant DMPs with the blue-red color scale representing the hypo- to hypermethylation trend. Hierarchical clustering of rows and columns represents beta-value of each group, respectively (p-value < 0.01). **d,** Gene ontology (GO) term enrichment analysis in wt, IDH1, KRAS, and TP53 mutation NPCs at 2 to 10 days comparisons (p-value < 0.01). GO terms show gene enrichment of wt and mutants are different, but three mutants share similarity. **e,** Venn diagram representing number of genes found within DMRs in IDH1, KRAS, and TP53 mutation cells compared with wt cells at 2 to 10 days and 10 to 10 + 2 iN days comparisons. **f,** Heatmap showing genes found within DMRs in IDH1, KRAS, and TP53 mutation cells compared with wt cells at 2 to 10 days and 10 to 10 + 2 iN days comparisons (p-value < 0.01). Color indicates maximum CpG difference within the DMR that the gene was found within. Geneset classification was identified using the hallmark genesets in GSEA. The gene classification is based on their major function and hallmark pathway signaling. **g,** Methylation beta-values across DMR where genes that flipped from hypo- to hypermethylation across time points were found. LIN28A in TP53 mutation cells, and FOXD3 in KRAS and IDH1 mutation cells. Asterisks represent DNA methylation marks at gene loci. **h,** Methylation beta-values across DMR where genes were found to be changed across mutational states (BAI1 and KMO) of IDH1, TP53, and KRAS mutation cells. Asterisks represent DNA methylation marks at gene loci.

At later time points we observed an increase in epigenetic variability within the mutant cell lines, notably for IDH1, KRAS, and Sox17 mutations (Fig. 3b). To measure oncogene-specific methylation changes, we identified differentially hyper-(beta-value > 0.8 and logFC > 0.3) and hypo-(beta-value < 0.2, logFC < -0.3) methylated CpG sites. We found that oncogenic mutation is associated with DNA methylation alteration in NPCs and NPC-derived iN cells generally (Extended Data Fig. 6a, b, c). Notably, the DNA methylation change in IDH1, TP53, and KRAS mutation NPCs are more significant, and these mutations are very common in brain tumors and pan cancer. Therefore, to further understand the properties of IDH1, TP53, and KRAS mutation, we identified DMPs at 2 days (Extended Data Fig. 6a, d), 10 days (Extended Data Fig. 6b, e) and 10 + 2 iN days (Extended Data Fig. 6c, f). The heatmap of significant DMPs demonstrated that the epigenetic perturbations were enhanced as induction time increased and differentiation occurred (p-value < 0.01) (Fig. 3c). To identify functional categories of differentially hypermethylated genes, we performed GO enrichment analysis using clusterProfiler, using a cut-off of 1.5-fold 5mC level change to define hypermethylated genes (meandiff > 0.05, padj < 0.01) (Fig. 3d). Genes are hypermethylated in wt NPCs at 2 to 10 days comparisons that are enriched among neural differentiation, regulation of neural precursor cell proliferation, endoderm development, stem cell differentiation, and cell fate commitment. On the contrary, genes are hypermethylated in IDH1, KRAS, and TP53 mutation NPCs at 2 to 10 days comparisons that are involved in the regulation of neuron death, neurotransmitter transport, cell proliferation, regulation of secretory pathway, endocrine system development, regulation of cation transmembrane transport, regulation of cell shape and growth. It was shown that there is a significant difference in gene methylation states between wt and mutants from 2 to 10 days. In addition, the three mutants share a number of hypermethylated genes.

To eliminate the effect that is not related with oncogenes, we did comparative analysis between wt-2days and wt-10days (wt-2/10days), between mutation-2days and mutation-10days (mutation-2/10days), then compared the wt-2/10days and mutation-2/10days. Similarly, we performed the comparison between wt-10days and wt-10+2iNdays (wt-10/10+2iNdays), between mutation-10days and mutation-10+2iNdays (mutation-10/10+2iNdays), then compared the wt-10/10+2iNdays and mutation-10/10+2iNdays (Extended Data Fig. 6g, h, Fig. 3e, f, g, h). Genes that were found to be differentially methylated in wt cells at the relevant comparison periods were removed in the following analysis. Notably, there are a large number of hypomethylated regions in mutant NPCs compared to wt NPCs from 2 to 10 days (Extended Data Fig. 6g, h). However, from 10 to 10 + 2 iN differentiation days, there are a large number of hypermethylated regions in mutant NPC-derived iNs compared to wt NPC-derived iNs (p-val < 0.05, max difference of CpG > 0.2) (Extended Data Fig. 6g, h). The distribution of DNA methylation change showed that most occurred at transcription start site ^9^ (Extended Data Fig. 6h).

To reveal the most distinct methylation changed genes, we delineated overlapping methylated genes between the three mutants at 2 to 10 days and 10 to 10 + 2 iN days comparisons (Fig. 3e). Interestingly, cells have IDH1, KRAS, and TP53 mutation share 25 genes that were hypomethylated at 2 to 10 days comparisons, and share 32 genes were hypermethylated at 10 to 10 + 2 iN days comparisons (Fig. 3e). Simultaneously, each mutant displays different methylation changed genes in a specific manner (Fig. 3e). We next categorized the methylation changed genes based on functionality (Fig. 3f). At 2 to 10 days comparisons, we found that NPCs with IDH1, KRAS, or TP53 mutation shared several of the same hypomethylated genes in the categories of pluripotency, multipotency, differentiation (BAI1, MTL5); neural development (CNPY1, GALR1, GBX2, LHX5); cancer (CACNA2D3, DMTN, FUCA2, IL12RB1, ITGAM, MYOZ3, NR2F1, NR2F1-AS1, UBR3, HIST1H1B) (Fig. 3f). At 10 to 10 + 2 iN days comparisons, we found that NPC-derived iN cells have IDH1, KRAS, and TP53 mutation shared several of the same hypermethylated genes in pluripotency, multipotency, differentiation (FOXD3, TAF10); neural development (GABRA1, GPR176, SALL1); cancer (FH, BAZ1A, DCLRE1C, FARSB, HOXC12, ILK, LMBR1, TPP1, UBE2T, ZNF540, ZNF571) (Fig. 3f).

In addition to the common methylation change in the three mutants, they also have distinct patterns. Hoxb1 is hypomethylated in IDH1 mutation NPCs, and KLF4 is hypomethylated in KRAS and TP53 mutation NPCs at 2 to 10 days comparisons (Fig. 3f). As reported, when Hoxb1 (Homeobox B1) expression is induced in ESC-derived NSCs (neural stem cells), few cells differentiate, but they keep proliferating ^10^. KLF4 (Kruppel Like Factor 4) -overexpressing NSCs resisted neuronal differentiation, showing rounded morphology with very short processes^11^, which is consistent with our observation. Additionally, at 10 to 10 + 2 iN days comparisons, Foxp1 is hypermethylated in IDH1 and KRAS mutation NPC-derived iN cells (Fig. 3f). Gain or loss of Foxp1 (Forkhead Box P1) function alters the level of progenitor cell divisions and the balance between self-renewal and differentiation^12^. The DNA methylation alterations happened mostly in the functional category related to multipotency and neural identification as well as tumorigenesis. Correspondingly, as previously mentioned, mutant stem cells resist neural differentiation and reprogramming as a consequence. DNA methylation changes in normal NPCs dynamically regulate cell fate and identity. However, aberrant DNA methylation changes in mutant NPCs dysregulate cell fate and identity.

To explore the critical effect of differentiation that occurs in mutant stem cells, we plotted the methylation of each DMP for a given region before and after differentiation (Fig. 3g). Interestingly, we found some genes were hypermethylated from 2 to 10 days, but convert to hypomethylated from 10 to 10 + 2 iN days, such as Lin28A in TP53 mutation cells (Fig. 3g). On the contrary, some genes were hypomethylated from 2 to 10 days, but convert to hypermethylated from 10 to 10 + 2 iN days, such as FOXD3 in IDH1 and KRAS mutation cells (Fig. 3g). Lin28A (Lin-28 Homolog A) controls NPC proliferation and neurogenesis in a SOX2-dependent manner^13^. Foxd3 (forkhead box D3) is associated with inactivation of important naive pluripotency genes by its modification of chromatin structures via recruiting histone demethylases and decreasing numerous activating factors^14^. These genes effect stem cell identification and differentiation regulation that were changed on their methylation modification in an aberrant manner, which severely effects the accuracy of lineage commitment.

Meanwhile, we found many overlapping gene sites were hypermethylated or hypomethylated in three mutants (Fig. 3f). In particular, BAI1 was hypomethylated from the 2 to 10 days, and KMO was hypermethylated from the 10 to 10 + 2 iN days (Fig. 3h). BAI1 (brain-specific angiogenesis inhibitor 1) is expressed in normal human brain, mediates p53 polyubiquitination^15,16^. KMO (Kynurenine 3-monooxygenase) influences stem cell biology and modulates neurodegeneration^17^. The three mutants share numerous genes with the same methylation site modification, which suggested that there is a specific methylation pattern generally related to oncogenic mutation associated with lineage differentiation disorder.

### Stem cells with oncogenic mutations share similar epigenetic patterns with brain tumors

As previously discussed, these oncogenes came from tumor cells via TCGA data analysis (Extended Data Fig. 1d), that are associated with DNA methylation alteration. It is important to evaluate the similarity of epigenetic states between mutant cells and cancer, such as low-grade gliomas (LGGs). To identify the methylation changed genes associated with LGGs, we analyzed the epigenetic profile of LGGs with IDH1-R132H, TP53-R273C, BRAF-V600E, or other mutations separately, from TCGA datasets as measured by DNA methylation, and compared to the non-tumor human whole brain cells as control^18^ (see Methods). To evaluate the similarity between mutant cells and LGG cells in DNA methylation patterns broadly, we compared the genomic regions that were differentially methylated between the control brain and LGGs with oncogenes, as well as between wt and mutant cells at 2, 10, and 10 + 2 iN days (Fig. 4a, b, Extended Data Fig. 7a, b). It was shown that many overlapping genes were hypermethylated in mutant cells and LGGs, and gene numbers increased as the increased days of oncogene induction and differentiation initiation (Fig. 4a, b), also many overlapping genes were hypomethylated (Extended Data Fig. 7a, b). The heatmap displays the most significantly differentially hypermethylated genes overlapped in mutant cells and cancer cells (Fig. 4b). Many genes retain the same methylation states from 10 to 10 + 2 iN days (Fig. 4b).

**Fig. 4.**
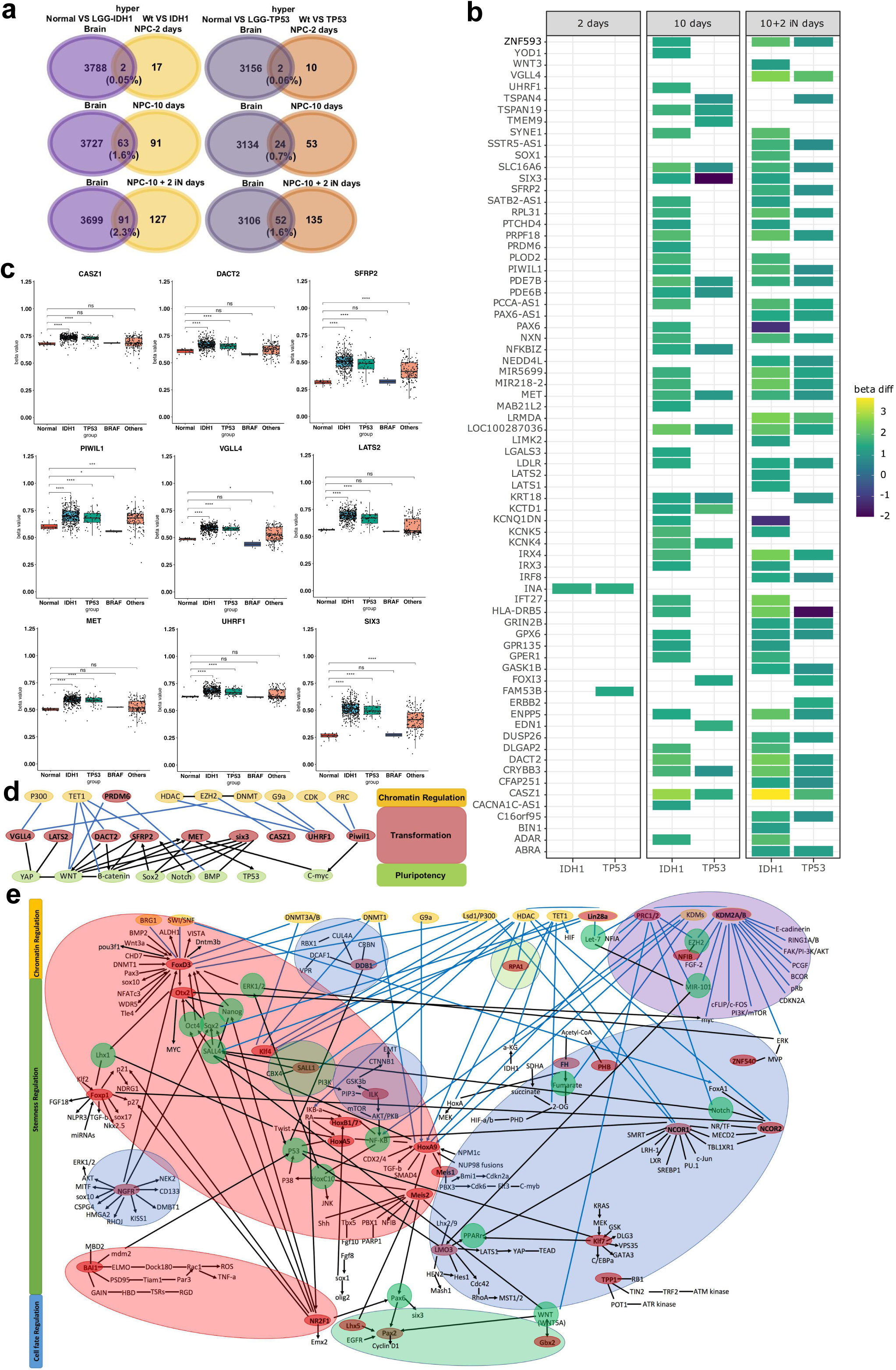
Stem cells with oncogenes share similar epigenetic patterns with brain tumors. **a,** Venn diagram representing number of hypermethylated genes found in DMRs come from the control brain VS low-grade gliomas (LGGs) with IDH1 or TP53 mutation, and wt cells VS IDH1 or TP53 mutation cells at 2, 10, and 10 + 2 iN days (p-value < 0.01). **b,** Heatmap of differentially hypermethylated genes found within the overlap node from Fig. 4a (p-value < 0.01). The colour scale indicates methylation level, from high (yellow) to low (blue). **c,** The box plot demonstrates the mean DNA methylation level and variation in methylation levels of genes from Fig. 4b in LGGs with IDH1-R132H, TP53-R273C, BRAF-V600E, other mutations and control brain cells as measured by DNA methylation from TCGA dataset (NS: p > 0.05, *: p <= 0.05, **: p <= 0.01, ***: p <= 0.001, ****: p <= 0.0001). The horizontal line in box-plots shows median values, and the lower and upper edges of the boxes represents the 25th and 75th percentiles, respectively, and upper and lower lines outside the boxes represent minimum and maximum values (error bars). Each black dot represents mean DNA methylation level of one affected case in the corresponding cohort. The methylation data of control brain came from whole human brain, and more details are in the method. **d,** Visualization of network of hypermethylated genes. The red circles mark genes from Fig. 4b, c. The yellow circles mark chromatin regulation enzymes are on the top. The green circles mark genes which play a critical role in pluripotency that connect with genes that are hypermethylated in LGGs and connect with genes which function as chromatin regulators. **e,** Visualization of the gene regulatory network. Biological Process categories were visualized as nodes clustered into functional modules and interconnected based on the indicated interaction confirmed in literature. There are five categories cycled as different colors: the red node indicates genes of pluripotency, multipotency, and differentiation (LIN28A, FOXD3, KLF4, HOXB1, MEIS, NR2F1, BAI1, HOXA5, HOXA9, PHB, FOXP1, OTX2); the green node indicates genes of neural generation and development (GBX2, LHX5, SALL1, PAX2); the blue node indicates genes of cancer (NCOR, NGFR, DDB1, FH, ILK, TPP1, ZNF540, LMO3, KLF7, NFIB); the light green node indicates genes that function in DNA, RNA binding and translation (RPA1); the purple node indicates genes that function in histone and chromatin regulation (KDM2A, PRC1/2). The red circles mark genes that are found to have methylation changes in Fig. 3f. The yellow circles mark chromatin regulation enzymes are on the top. The green circles mark genes that connect with at least three other genes. The black lines represent the interaction of two genes which has been confirmed in literature. The blue lines represent the genes connected with chromatin-modifying regulators. The arrows indicate an effect of one gene on another gene.

We found that many genes were hypermethylated in NPCs and NPC-derived iN cells with oncogenes, that were also hypermethylated in LGGs with the same oncogene (Fig. 4c), such as CASZ1, DACT2, SFRP2, MET, PIWIL1, VGLL4, LATS2, SIX3, UHRF1. As reported, SFRP2 is downregulated in glioma patients by promoter hypermethylation, and regulates Wnt/β-catenin activation in glioma cells^19,20^. Inactivation of LATS1/2 kinases causes YAP/TAZ-driven global hypertranscription in neural progenitors that inhibits differentiation^21^. LATS1/2 deficiency reprograms schwann cells (SCs) to a cancerous, progenitor-like phenotype and promotes hyperproliferation^22^. Notably, to fully understand the relation of these hypermethylated genes, we analyzed the biological connection of them. We found that the hypermethylated genes in both mutant cells and tumor cells have a close relation to chromatin-modifying enzymes (Fig. 4d).

Moreover, we compared the genomic regions that were differentially methylated between the control brain and LGGs with oncogenes, as well as between wt and mutant NPCs or NPC-derived iN cells at 2 to 10 days or 10 to 10 + 2 iN days comparisons (Extended Data Fig. 7c, d). Notably, we found that the genes were hypermethylated in mutant cells from Fig. 3f and were also hypermethylated in LGGs (Extended Data Fig. 7c, d), such as TMEM106A and RARRES1 at 2 to 10 days comparisons (Extended Data Fig. 7e), Foxd3 and HOXC12 at 10 to 10 + 2 iN days comparisons (Extended Data Fig. 7f). These genes were hypermethylated in mutant cells and LGGs that have a critical function in cell fate determination via epigenetic modification. Collectively, tumors with various accumulated oncogenes share similar DNA methylation patterns with stem cells have oncogenes after a short period of induced expression.

To specifically determine DNA hypermethylation sites consequent to oncogenic mutation associated differentiation impairment, we investigated each methylation locus of hypermethylated genes from Fig. 3g, h, 4c in mutant cells and brains. Elevated methylation occurred at the same loci of genes in mutant cells and LGGs, such as CASZ1 and VGLL4 (Extended Data Fig. 7g).

Furthermore, we analyzed the biological connection of key genes from Fig. 3f, in order to show the whole picture of the network. Biological process categories were visualized as nodes clustered into functional modules and interconnected based on the indicated interaction confirmed in literature. The red circle labeled genes had methylation alteration as shown in Fig. 3f. We categorized the genes by five different colors: the red node indicates genes in pluripotency, multipotency, and differentiation (LIN28A, FOXD3, KLF4, HOXB1, MEIS, NR2F1, BAI1, HOXA5, HOXA9, PHB, FOXP1, OTX2), the green node indicates genes in neural development (GBX2, LHX5, SALL1, PAX2), the blue node indicates genes in cancer (NCOR, NGFR, DDB1, FH, ILK, TPP1, ZNF540, LMO3, KLF7, NFIB), the light green node indicates genes in DNA, RNA binding and translation (RPA1), the purple node indicates genes which function as histone and chromatin enzymes (KDM2A, PRC1/2) (Fig. 4e). The chromatin-modifying enzymes are labeled in yellow circles and shown on top of the visualization. We labeled genes connected with at least three other genes with green circles. The interaction of genes was connected with black lines based on published literature. We applied blue lines to indicate genes connected with chromatin-modifying regulators. The arrow indicates the effect of one gene on another gene. The gene regulatory network demonstrated that the genes play a critical role in multipotency maintenance, lineage differentiation, and cancer formation connect with each other via epigenetic regulation of chromatin-modifying enzymes.

To summarize the findings from this comprehensive data analysis, it is not only IDH1 mutation that causes methylation change ^3,4^, but other mutations, such as KRAS, TP53, and SOX17 mutation. The DNA methylation states are important to the maintenance of specific cell types, and also to facilitate differentiation from one cell type to another. Oncogenic mutations occurred in stem cells and progenitor cells which perturb DNA methylation states on lineage commitment genes and their functional performance in the process of differentiation, consequently, emerging a type of cell that maintain multipotency, resist differentiation, and retain proliferation, that is a novel exploration of impairment of differentiation efficiency. We are the first to identify oncogenic mutations that disrupt DNA methylation change on lineage differentiation genes, therefore, mutant stem cells undergoing inaccurate lineage commitment.

## Discussion

In 2017, Chao et al. generated iPSCs from AML patient samples and found that AML-iPSCs lacked leukemic potential, but when differentiated into hematopoietic cells, they reacquired the ability to give rise to leukemia *in vivo* and reestablished leukemic DNA methylation/gene expression patterns^1^. To investigate the relationship of oncogenic mutation and DNA methylation change during lineage differentiation, we constructed mutant NPCs (or hESCs) for comparing the lineage differentiation efficiency with or without oncogenes. We found stem cells with oncogenes were hard to reprogram and differentiate. During reprogramming and differentiation, a global reset of the epigenome occurs^8^. Also, the epigenetic states are more permissive for gene expression during differentiation. It implies the possibility that these stem cells with oncogenes may initiate differentiation failures via taking advantage of epigenetic reset during lineage differentiation.

To investigate the essential DNA methylation changes that occurred in mutant NPCs at the stage of maintenance and differentiation, we performed methylation assay on cells undergoing oncogene induction for 2 days or 10 days, and neural cell differentiation for 2 days. We found that Lin28A was hypermethylated in TP53 mutant cells at 2 to 10 days, but converts to hypomethylated at 10 to 10+2 iN days (Fig. 3g). In the opposite way, some genes were hypomethylated at 2 to 10 days, but convert to hypermethylated at 10 to 10+2 iN days, such as FOXD3 in IDH1 and KRAS mutant cells (Fig. 3g). It had been reported that Lin28 regulates the self-renewal of stem cells through direct interaction with target mRNAs and by disrupting the maturation of certain miRNAs involved in embryonic development^13^. Lin28a/b double knockout mice display neural tube defects and reduced proliferation and precocious differentiation of NPCs^23^. FOXD3 was demonstrated to be required in maintaining pluripotency in mouse embryonic stem cells^14^. Importantly, the DNA methylation states of these genes are converted in mutant NPCs compared to wt NPCs during lineage differentiation. The mechanism of converted methylation states corresponding with differentiation is interesting to explore further.

As found previously, these oncogenes came from tumor cells as shown in Extended Data Fig. 1d. Therefore, we evaluated the similarity of epigenetic states between mutant cells and LGGs. Importantly, it was shown that many genes that were hypermethylated in mutant cells also were hypermethylated in LGGs. CASZ1 functions as a neural fate determination gene and tumor suppressor. Piwil1 (piwi like RNA-mediated gene silencing 1) plays an important role in RNA silencing and stem cell self-renewal^24^. Six3 (SIX homeobox 3) is required for development of the rostral forebrain through binding to the Wnt1 promoter region^25^. Uhrf1 (ubiquitin like with PHD and ring finger domains 1) is a widely known hemi-methylated DNA-binding protein, playing a role in DNA methylation through the recruitment of Dnmt1^26^. Uhrf1 regulates cell specification, particularly those associated with neuroectoderm and mesoderm specification^27,28^. SFRP2 (secreted frizzled related protein 2) is a Wnt antagonist and inhibits β-catenin^29^ and bone morphogenic protein (BMP) signaling pathways, and SOX2 directly binds the SFRP2 promoters^30^. SFRP2 improves mesenchymal stem cells engraftment by modulating self-renewal through increasing stem cell survival and by inhibiting differentiation^19^. The inactivation of Hippo pathway kinases LATS1/2 (large tumor suppressor kinase 1/2) causes a YAP/TAZ-driven global increase in transcription activity during brain development, and upregulates many genes involved in cell growth and proliferation, resulting in inhibition of differentiation of neural progenitor cells ^21^.

In summary, our findings reveal an essential function of these lineage differentiation genes as a stabilizer of the epigenome by promoting epigenetic modifications which are necessary for the faithful differentiation and the maintenance of pluripotency and multipotency (Fig. 4e). There is a lineage-specific epigenetic pattern. When stem cells have oncogenes, oncogenic mutation associated epigenetic change effects the expression of pluripotent genes, and impair the accuracy of lineage commitment.

## Methods

### Key resources table

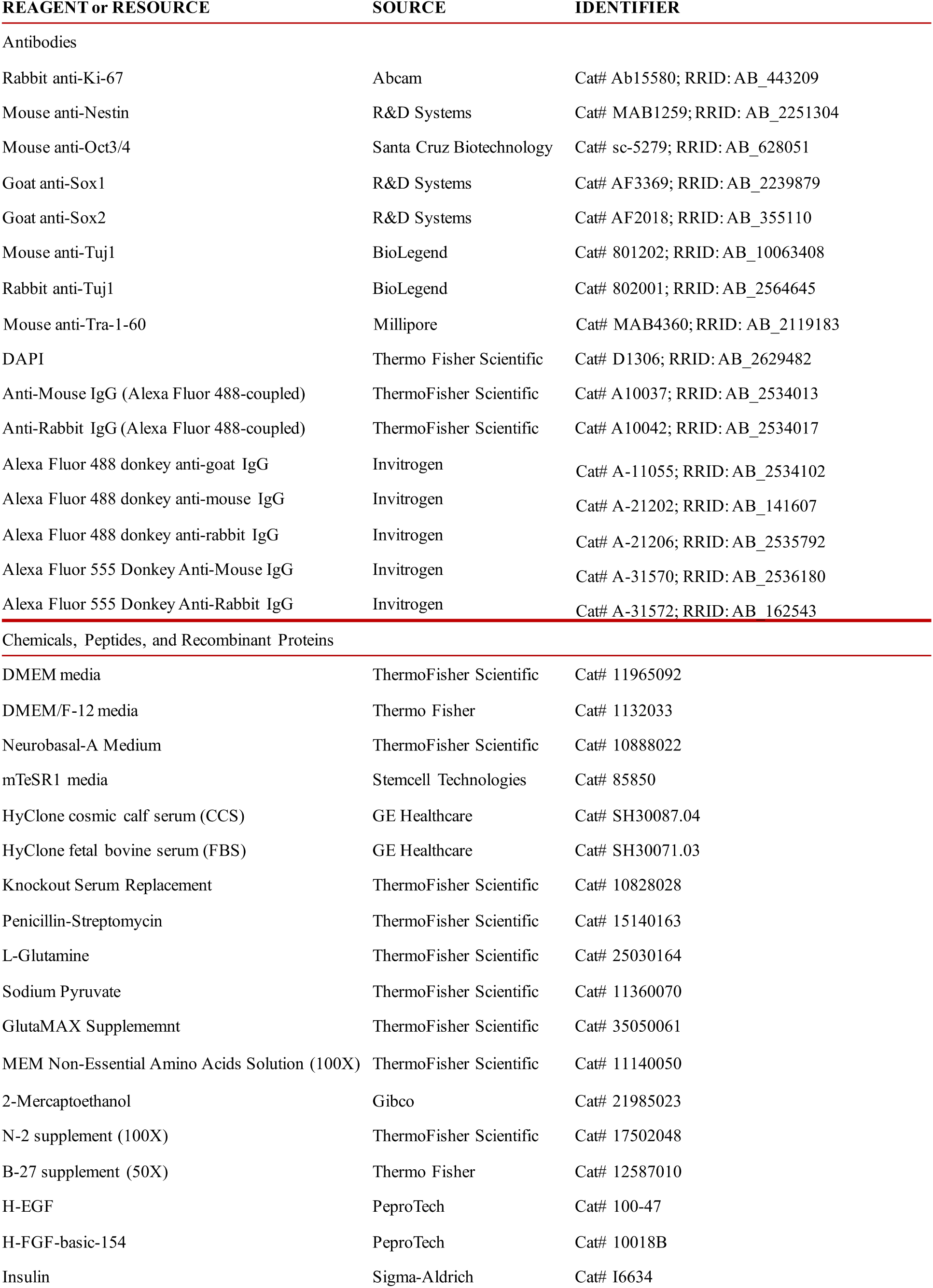
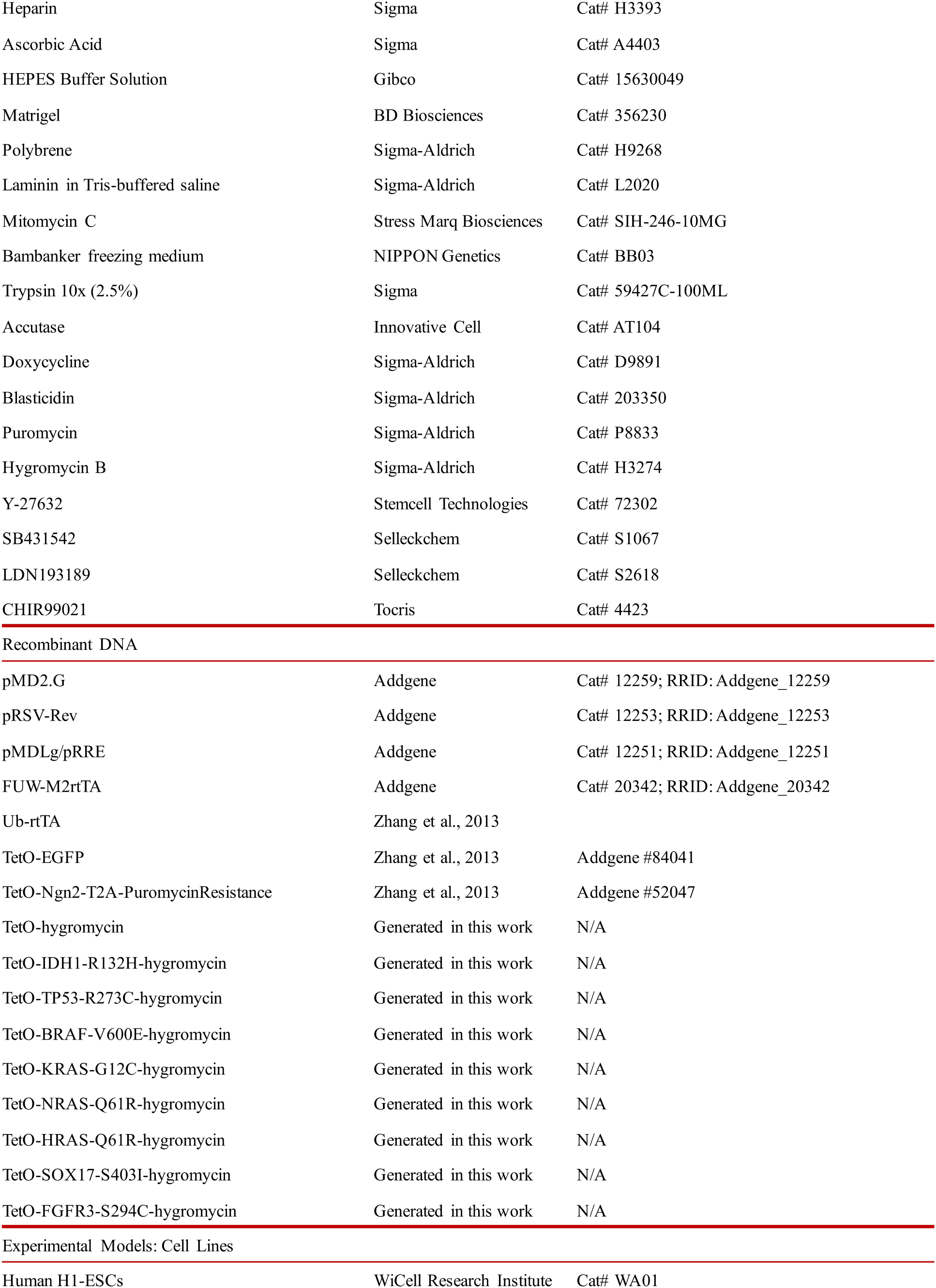
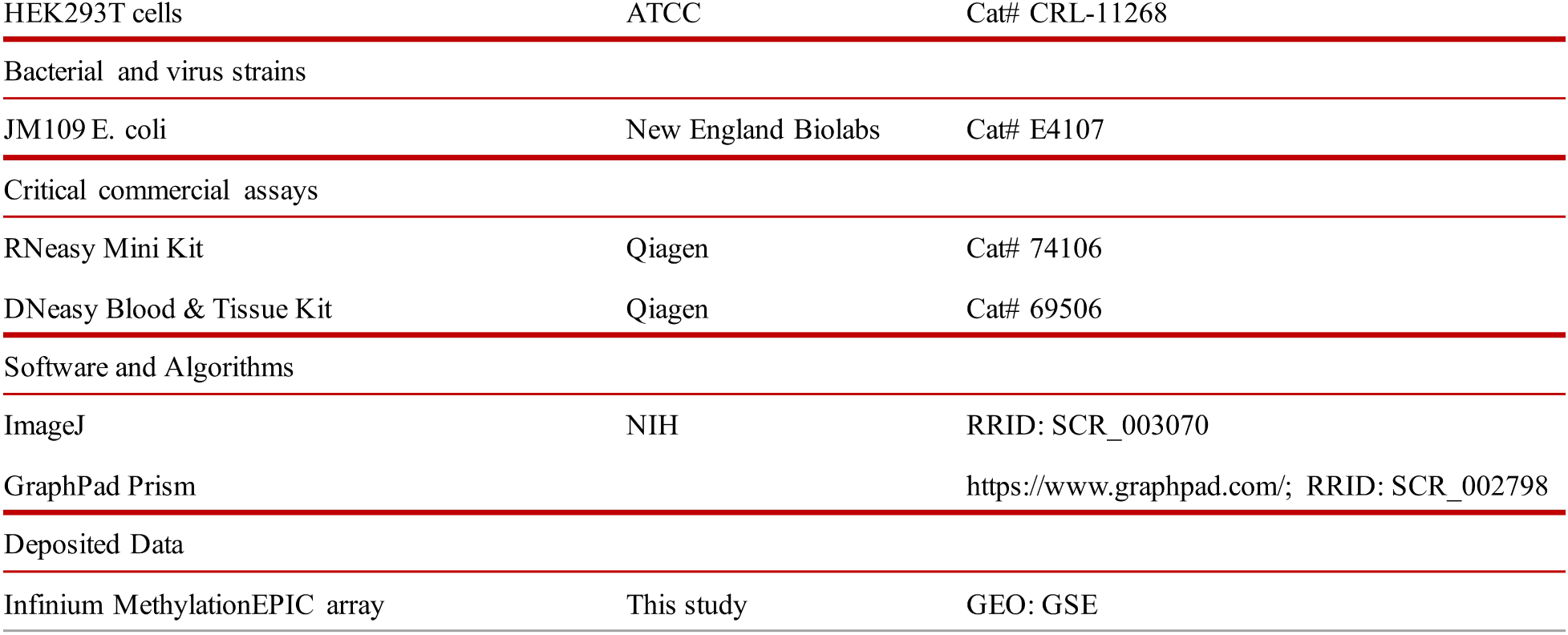

### Cell culture and media

H1 ES cells were obtained from WiCell Research Resources (WiCell, WI). ES cells were maintained as feeder-free cells in mTeSR™1 medium (Stem Cell Technologies). We used a doxycycline (dox)-inducible lentiviral vector system to transduce empty vector plasmid as a control and oncogenes into the H1 ES cells, separately. Selection was carried out for 7 days in mTeSR™1 medium with 50 μg/ml hygromycin.

### Generation of iN cells from human ES cells

H1 human ESCs were dissociated using accutase and plated as single cell in mTeSR™1 medium supplemented with ROCK inhibitor Y-27632 (10 μM, StemCell Technologies, Inc.) on matrigel to a final density of 20,000-30,000 cells per cm^2^. On the second day, attached cells were infected with lentivirus for Ngn2-puro and rtTA expression. After 24 hr (day 0), transgene expression was induced by replacing the medium with N2/B27 medium (DMEM/F12, 1% N2 supplement (Life Technologies), 1% B27 supplement (Life Technologies), 10 μg/ml insulin (Life Technologies)), and 2 μg/ml Doxycycline. Selection was carried out for 14 days (day 1 to day 14) in N2/B27 medium with 2 μg/ml puromycin. From day 2, induced neuronal (iN) cells were maintained in N2/B27 medium and changed medium every 2 days. Neuronal cells were defined as cells that stained positive for Tuj1 and had a process at least three times longer than the cell body.

### hESCs differentiate to neural progenitor cells

hESCs were grown in mTeSR media on matrigel coated dishes until confluent, then enzymatically passaged with accutase, resuspended in Neurobasal-A medium, DMEM/F12, 1% N2, 1% B27 and insulin media, with Y-27632, a specific inhibitor of Rho kinase (ROCK) activity, and cultured in the same size of the dish without matrigel coating. Overnight, colonies will form floating spherical clusters termed “embryoid bodies” (EBs). When EB size grows to 200 ~ 400 μm, they were gently moved to matrigel coated dishes. The next day, after the EBs attached to the bottom, the media was changed to N2/B27 media supplemented with dual SMAD inhibitors (LDN193189 and SB431542). EBs were fed every second day with N2/B27 media supplemented with 0.1 μM LDN193189 and 10 μM SB431542^7^. Cells are cultured for several days, until neural rosettes begin to appear, characterized as round clusters of neuroepithelial cells with apicobasal polarity. Staining of Sox1 is performed to identify the neural progenitor cells.

### Optimized protocol for NPCs differentiation from hESCs

NPCs were differentiated from hESCs using dual SMAD inhibitors^7^ (Extended Data Fig. 3). High-density hESCs were identified on day 1, and passaged to the same size of dish to form embryoid bodies in iN cell medium on day 2~3 until the EB size is large as 200~300 μm. EBs were harvested and moved to matrigel coated dishes on day 4. Media was changed to NSC medium with TGF-β and BMP inhibitor to differentiate NPCs from day 5 to day 21. NSC basal medium includes DMEM/F12 and Neurobasal medium at 1:1, 1% Sodium pyruvate, 1% P/S, 1% NEAA, 1 M HEPES, 1% Glutamax, 1% NaHCO3. When used for culturing cells add: 1% B27, 20 ng/ml bFGF, 20 ng/ml EGF, 1 μl/ml B27 supplement, 2 μg/ml Heparin, and 0.1% LIF. NPC differentiation medium includes fetal NSC basal medium, 1% N2, 2% B27, 0.1 mM Ascorbic acid, 10 μM SB431542 (TGF-β inhibitor), and 0.1 μM LDN193189 (BMP Inhibitor)^31^. The neural rosette formed around day 6. Cells were passaged and medium changed to NSC medium on day 22. Immunostaining for the pluripotency marker Oct4 and NPC marker Sox1 for NPCs was performed on day 28 (Extended Data Fig. 3). As low density will cause NPC differentiation, the cells were plated at very high density. After four weeks of culturing in NSC medium with dual SMAD inhibitors, all the cells are Sox1 positive NPCs, and there are no Oct4 positive cells present in all field tested by immunofluorescence. However, not all the hESCs successfully differentiate into NPCs in the dual SMAD inhibitors NSC medium. Mesoderm and endoderm cells spontaneously showed in the process. NPCs come from the neural rosette, pick up the neural rosette when passage cells, and maintain NPCs in NSC medium monthly which could potentially help to purify NPCs.

### Neural progenitor cells reprogram to iPSCs

The dox-inducible lentiviral transduction of the transcription factors Oct4, Sox2, c-Myc, and Klf4 (OSKM) is an efficient method to reprogram human NPCs to iPSCs^32,33^. We transduced cells with dox-inducible lentiviral vectors carrying the mouse cDNAs encoding the four transcription factors Oct4, Sox2, c-Myc, and Klf4. These cells were simultaneously infected with a constitutively active lentivirus transducing the reverse tetracycline transactivator (FUW-M2rtTA). The infected cells were cultured in the presence of dox, and iPSCs with a human ESC like morphology were detected after approximately 2 weeks in four factors culture.

### Virus generation

Lentiviruses were produced as described^33^ in HEK293T cells (ATCC, VA) by co-transfection with three helper plasmids (pRSV-REV, pMDLg/pRRE and vesicular stomatitis virus G protein expression vector) (12 μg of lentiviral vector DNA and 6 μg of each of helper plasmid DNA per 75 cm^2^ culture area) using polyethylenimine (PEI) ^34^. Lentiviruses were harvested with the medium 24 and 46 hr after transfection. Viral supernatant was filtered through a 0.45 μm filter, pelleted by centrifugation (21,000 × g for 2 hours), resuspended in DMEM, aliquoted, and snap-frozen in liquid N_2_. Only virus preparations with >90% infection efficiency as assessed by EGFP expression or puromycin resistance were used for experiments.

### Lentiviral infections

Complementary DNAs for candidate genes were cloned into doxycycline-inducible lentiviral vectors, as described previously^35^. Lentiviral production and hESCs, NPCs infections were performed as described previously^36^. Human ESCs and NPCs were infected with concentrated lentivirus and treated with doxycycline (2 mg/μl) 16~24 hr later.

### Immunofluorescence

Cultured iN cells were fixed in 4% paraformaldehyde in PBS for 15 min at room temperature, washed three times with PBS, and incubated in 0.2% Triton X-100 in PBS for 10 min at room temperature. Cells were blocked in PBS containing 10% BSA for 3 hours at 4℃. Primary antibodies were applied overnight. Cells were washed in PBS three times. Secondary antibodies were applied for 3 hours and then washed. Cells were stained with 300 nM DAPI for indication of nuclei, and mounted with mounting medium (containing antifade agent). The coverslips were sealed, and the target antigen was visualized by fluorescence microscopy.

### Antibody

The following primary antibodies with indicated dilution in blocking buffer were used: Rabbit anti-Tuj1 (Covance, BioLegend, MRB-435P, 1:1,000), Mouse anti-Tuj1 (Covance, BioLegend, MMS-435P, 1:1,000), Rabbit anti-human Sox1 (Millipore, AB15766, 1:1000), Mouse anti-Tra-1-60 (Millipore, MAB4360, 1:1000), Rabbit anti-Ki67 (Abcam, ab15580, 1:1000), Goat anti-Sox2 (Santa Cruz, Y-17, sc-17320, 1:1000), Mouse anti-Sox2 (R&D, MAB2018, 1:200), Goat anti-Sox2 (R&D, AF2018, 1:200), Mouse anti-Oct-3/4 (Sata Cruz, C-10, sc-5279), Mouse anit-Nestin (DSHB, rat-401, 1:200), Mouse anit-Nestin (R&D, MAB1259, 1:200), Rabbit anti-ASCL1 (Abcam, ab74065, 1:200). After staining with corresponding secondary antibodies in PBS plus 1% BSA, coverslips were washed four times with PBS, each for 10 min, mounted with the mounting medium onto glass slides, and examined under Olympus FluoView FV1000.

### qRT-PCR

RNA was isolated through standard methods following manufacturer’s instructions (QIAGEN), and reverse transcribed with SuperScript III (Life Technologies). qRT-PCR was performed with FastStart Universal SYBR Green Master Mix (Roche) along with gene specific primers on a 7900HT real-time PCR machine (Applied Biosystems). Statistical significance was determined by Students t test based on triplicated experiments.

### DNA methylation analysis using Infinium MethylationEPIC array

Genomic DNA was isolated from the indicated hESC, NPC, and NPC-derived iN cell populations through standard methods (QIAGEN) according to the manufacturer’s instructions. Samples were processed and run at the Stanford Functional Genomics Facility (SFGF) using the Infinium MethylationEPIC array which interrogates over 850,000 methylation sites at single-nucleotide resolution. All data files that were used in our analysis can be found at NCBI. The GEO accession numbers are GSE268459, GSE268464, GSE268467, GSE268469, GSE268471. The control brain cells came from the whole human brain^18^.

### Calculating DMPs and DMRs across mutations and timepoints

The ChAMP (Chip analysis methylation pipeline) R package was used to normalize the dataset across all cell lines and time points using the BMIQ (Beta-mixture quantile dilation) normalization method. Each DMP and DMR calculation was done in a pairwise fashion for each mutation and wt cell line independently. We compared each mutant and wt cells at 2 days, 10 days, and 10 and 2 days of induction. In addition, 2 days vs. 10 days induction time points were compared together. 10 days vs. 10 days and 2 days of induction were compared together. DMPs were identified using the limma R package. Hypermethylated DMPs were determined as positions that had a beta-value greater than 0.85 and a log fold-change greater than 0.25. Hypomethylated DMPs were determined as positions that had a beta-value less than 0.25 and a log fold-change less than 0.35. At 2 to 10 days comparisons and 10 to 10 + 2 iN days comparisons, the same DMPs for the timeline of comparison calculated in wt cell lines were removed in order to identify mutation-specific DMPs. DMRs were calculated using the Bumphunter R package and annotated with the ilm10b4.hg19 genome. Both hypomethylated and hypermethylated DMRs were defined as regions with FDR < 0.1, number of cpgs < 2, and absolute maximum difference of CpG expression > 0.2. UCSC genome browser (GRCh37/hg19) and Integrative Genomics Viewer (IGV) illustration of the methylation regions were shown in Fig. 3, 4 and Extended Data Fig. 5, 6, 7. The GO analysis was performed with ClusterProfiler. The DNA methylation data included in this publication have been deposited in NCBI’s Gene Expression Omnibus repository (GSE). https://www.ncbi.nlm.nih.gov/geo/info/geo_illu.html

### Statistical Tests

Microsoft Excel was used for data analysis. Cumulative plots were generated from parameters collected from individual cells under similar experimental conditions. Average data are presented as bar graphs indicating means ± SD. Statistical comparisons between bar graphs were made using the unpaired, two-tailed, Student t test (NS: p > 0.05, *p < 0.01, all versus control).

## Acknowledgements

We thank the Stanford Functional Genomics Facility (SFGF) for providing excellent sequencing by using the Infinium MethylationEPIC array. We also would like to thank Daniel Haag, Qianyi Lee, Gernot Neumayer, Yohei Shibuya, Wendy Mei Fong, Nan Yang, Samuele Giuseppe Marro, Soham Chanda, Cheen Euong Ang, Bo Zhou, Bahareh Haddad Derafshi, Madhuri Vangipuram for critical discussions, sharing reagents, and providing technical assistance. Funding: This study was supported by grants from the NIH (R01) and Emerson Collective.

## Author Contributions

L.M. conceived and designed the study; L.M., A.Y., H.H., D.A.K., J.M., and Y.Y. planned and performed the experiments; A.Y., H.H., D.A.K., and J.M. performed DNA methylation data analyses; L.M. provided critical advice on experimental designs and data interpretations; K.S. and J.A.J provided experimental material; L.M. wrote the manuscript. L.M., K.S., A.Y., and D.A.K. edited the manuscript and the figures.

## SUPPLEMENTARY MATERIAL

**Extended Data Fig. 1.**
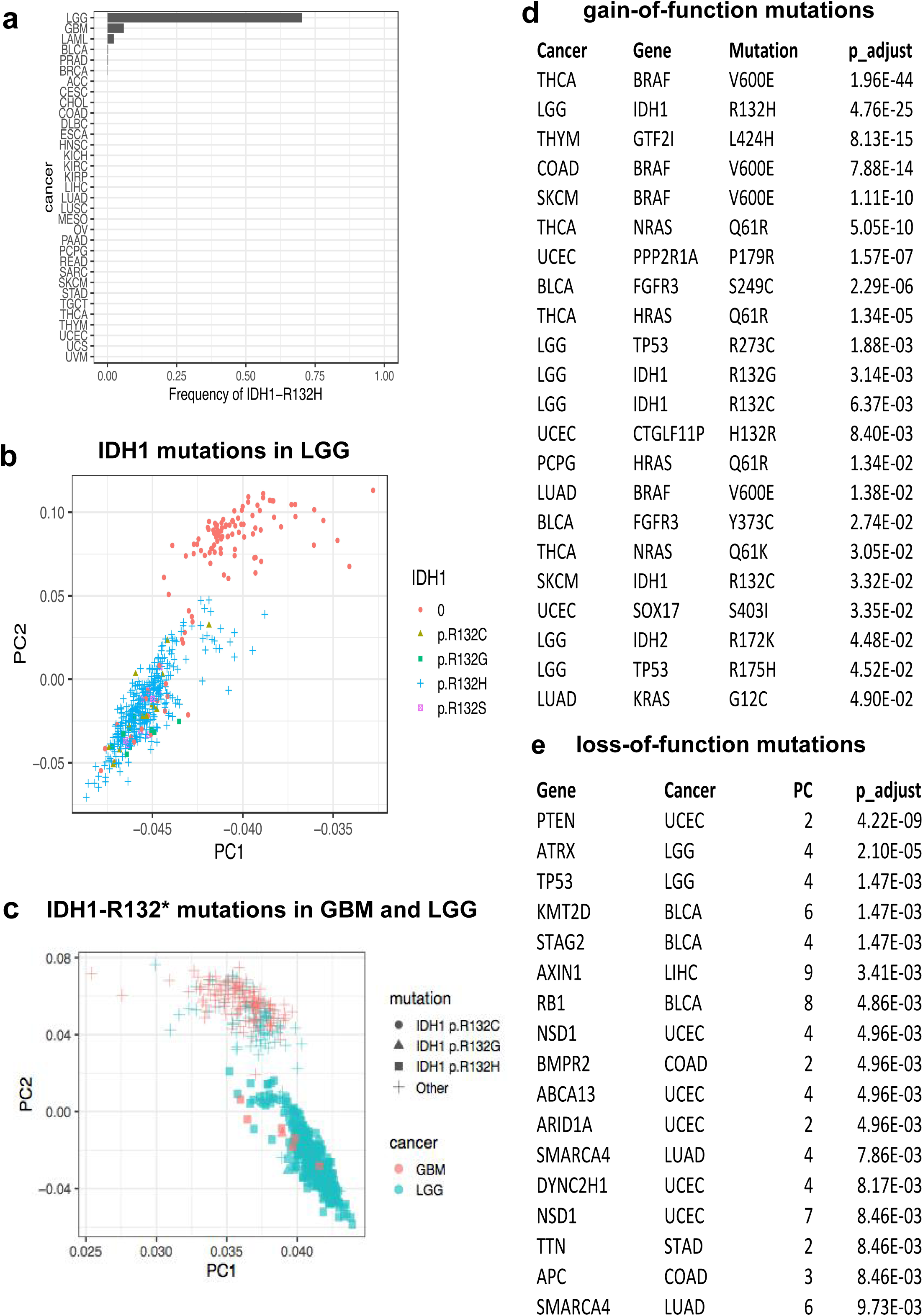
Oncogenic mutations are associated with epigenetic change from the The Cancer Genome Atlas dataset. **a,** Depicting the frequency of IDH1 mutation within each cancer type analysed. **b, c,** The effect of IDH1-R132* mutations on epigenetic state in low-grade glioma (LGG) and glioblastoma (GBM) as measured by DNA methylation. **d, e,** The 22 and 17 candidate genes in gain-of-function and loss-of-function mutations respectively (1% FDR, false discovery rate) involved in regulating the epigenetic landscape in cancer, by performing a cancer analysis linking both gain-of-function and loss-of-function mutations to changes in the epigenetic state by measure DNA methylation.

**Extended Data Fig. 2.**
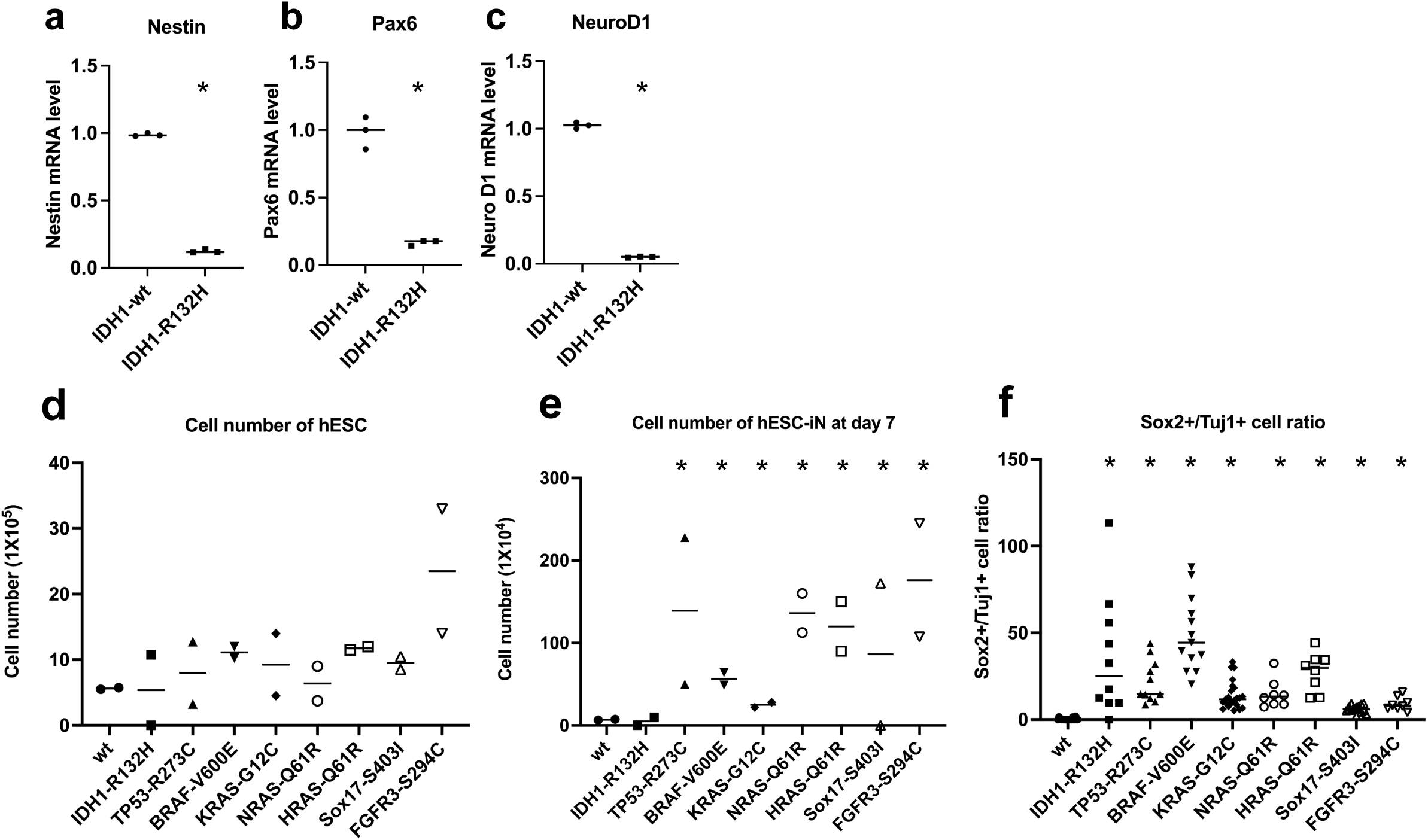
Stem cells with oncogenic mutations retain their potential for proliferation in differentiation. **a,** Cell number counting of wt hESCs and mutant hESCs after 3 days of dox-induction (n=3, NS: p > 0.05). **b,** Cell number counting of wt hESC-derived iN cells and mutant hESC-derived iN cells at 7 days of differentiation (n=3, *p < 0.01). **c,** The ratio of the Sox2 positive hESC-derived iN cell number and Tuj1 positive hESC-derived iN cell number at 10 days of differentiation (*p < 0.01).

**Extended Data Fig. 3.**
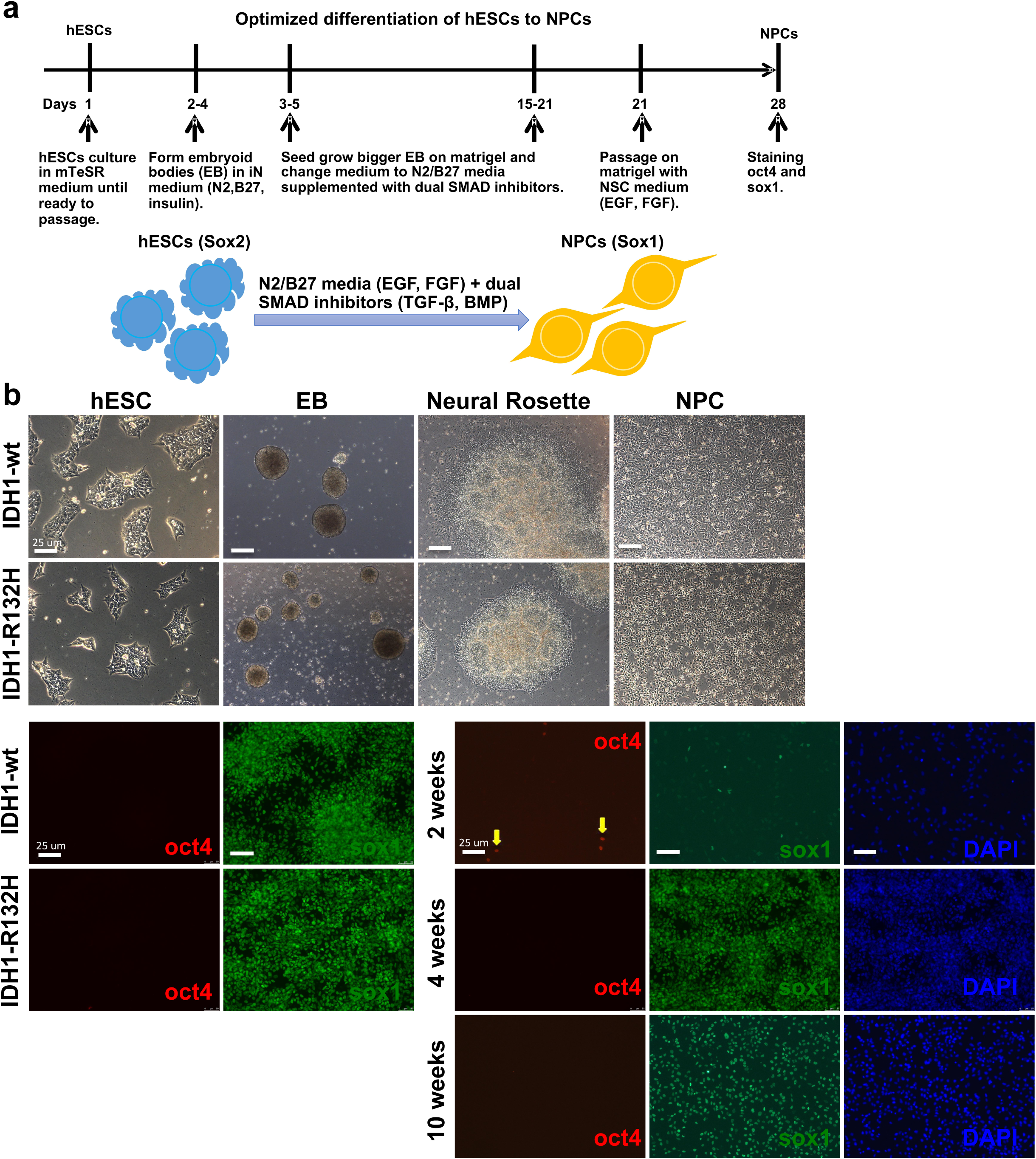
Optimize differentiation protocol for generation of NPCs from hESCs. **a,** Experimental scheme for generating human NPCs differentiated from hESCs use dual SMAD inhibitors. **b,** Brightfield of hESCs at day 1, embryoid bodies (EBs) at day 3, neural rosette at day 6, photo NPCs at day 28, and immunostaining pluripotency marker Oct4 (red) (yellow arrow) and NPC marker Sox1 (green) for NPCs at day 28. Nuclear staining by DAPI (blue). Scale bars, 25 μm.

**Extended Data Fig. 4.**
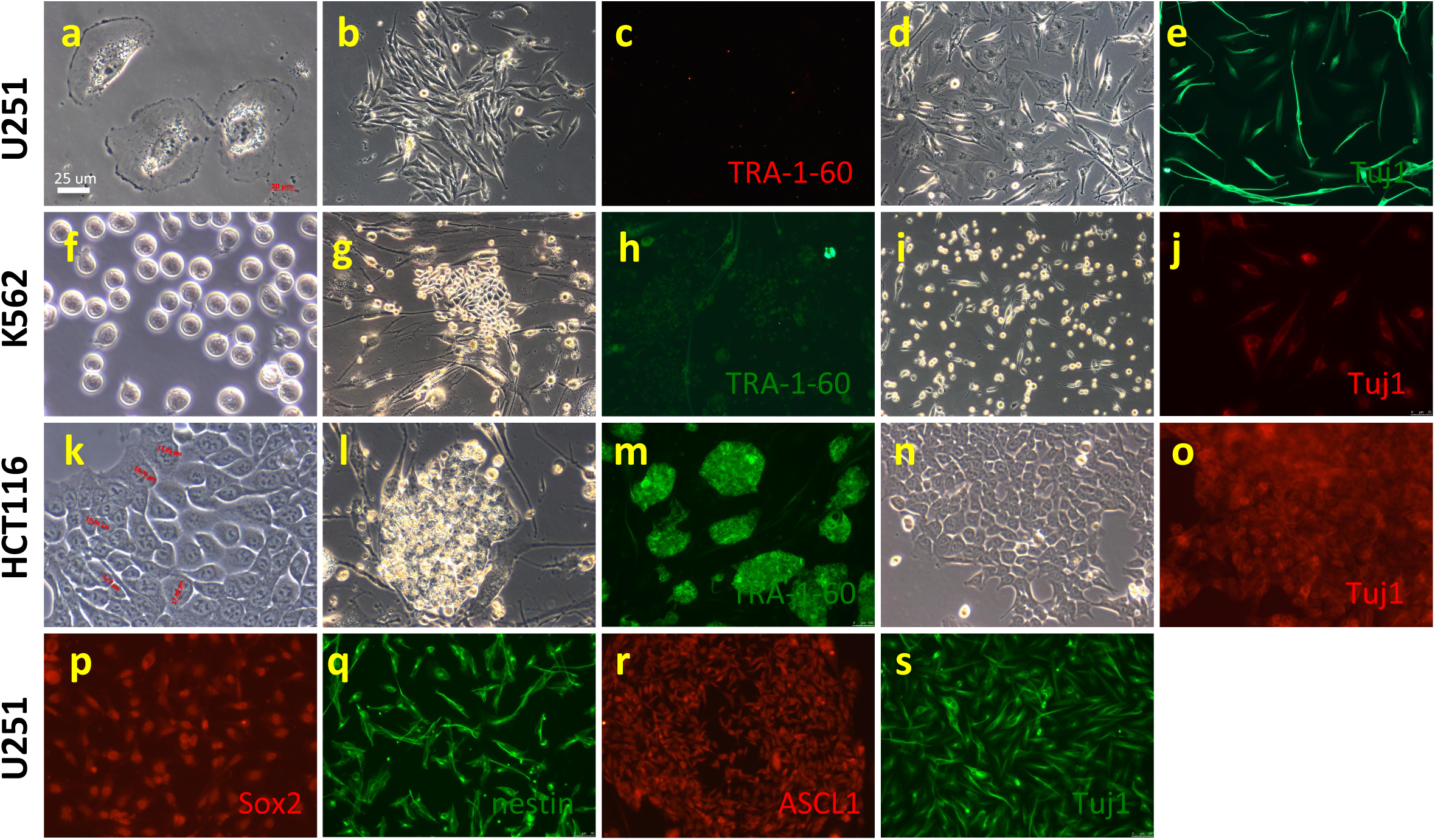
Cancer cells are difficult to reprogramming and differentiation. **a,** The morphology of U251 cells. U251 cell line is an IDH1 mutation glioblastoma cell line. Scale bars, 25 μm. **b, c,** Brightfield and fluorescent images of immunostaining TRA-1-60 (red) of U251 cells at 14 days reprogramming with OSKM for colony identification. **d, e,** Brightfield and fluorescent images of immunostaining U251 cells at 14 days differentiation by Ngn2 overexpression for the neural cells marker Tuj1 (green). **f,** The morphology of K562 cells. K562 cell line is an IDH1 mutant leukemia cell line. **g, h,** Brightfield and fluorescent images of K562 cells immunostaining for TRA-1-60 (green) at 14 days reprogramming with OSKM. No TRA-1-60 positive staining colony. **i, j,** Brightfield and fluorescent images of staining Tuj1 (red) and DAPI (blue) on K562 cells at 14 days differentiation by Ngn2 overexpression. **k,** The morphology of HCT116 cells. HCT116 cell line is an IDH1 mutant colon cancer cell line. **l, m,** Brightfield and fluorescent images of staining TRA-1-60 (green) and DAPI (blue) on HCT116 cells at 14 days reprogramming with OSKM. HCT116 cells cluster on feeders and grow more as colony-like clumps as shown in the photo. All cells are TRA-1-60 positive. **n, o,** Brightfield and fluorescent images of staining Tuj1 (red) and DAPI (blue) on HCT116 cells at 14 days differentiation by Ngn2 overexpression. **p, q, r, s,** Immunostaining U251 cells with the pluripotency marker Sox2 (red), neural stem/progenitor cell markers Nestin (green) and ASCL1 (red), and neural cell marker Tuj1 (green).

**Extended Data Fig. 5.**
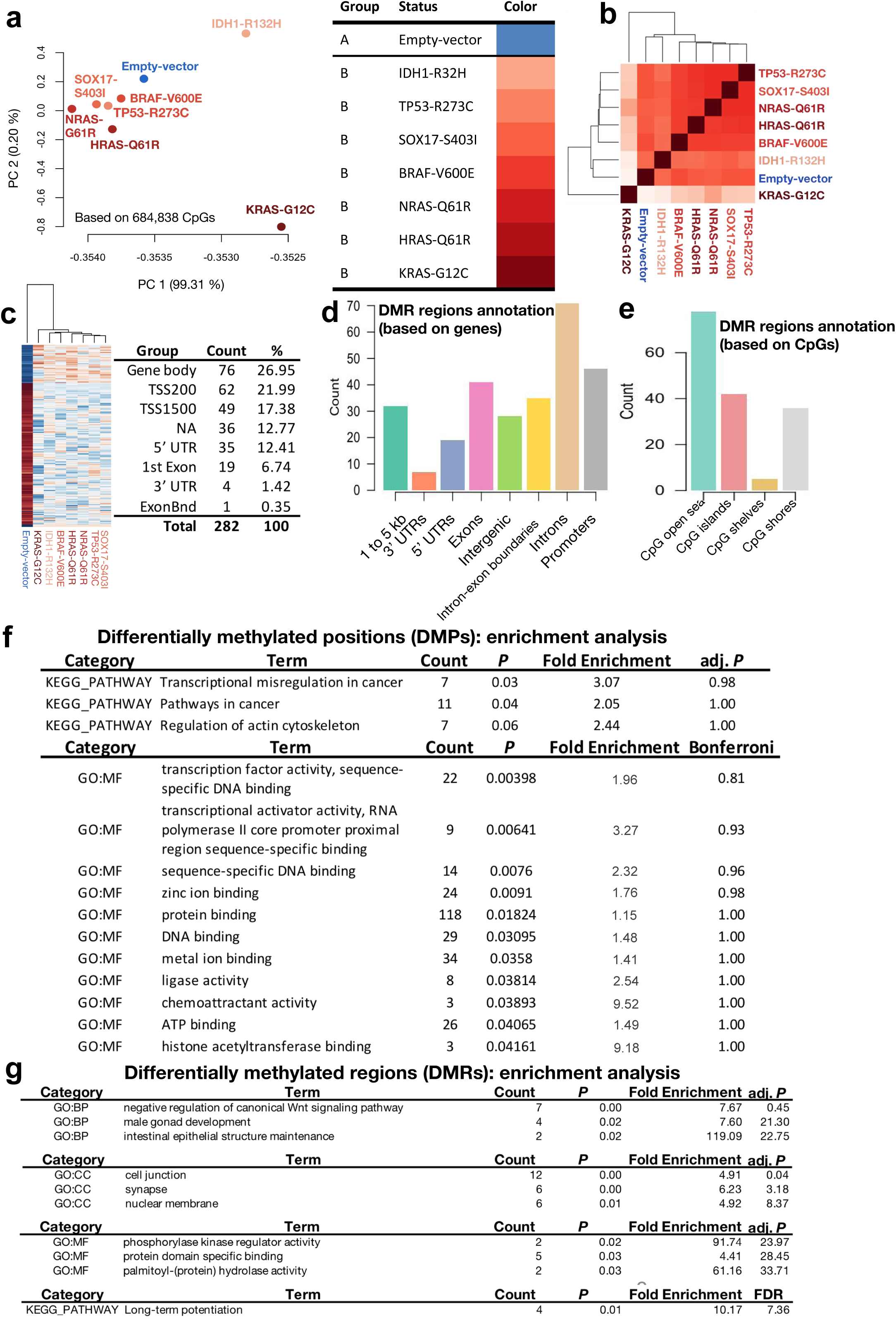
Oncogenic mutations are associated with epigenetic change in hESC cells. **a,** Principal component analysis (PCA) of DNA methylation profiles using Infinium MethylationEPIC BeadChip assay demonstrates that seven oncogene mutants and one wt hESC cells are distinguished by DNA methylation states. Each point represents wt as control or a gene modified cell sample. The proportions of variance explained by PC1 and PC2 are indicated. **b,** Heatmap plot shows the potential involvement of the most significant DNA methylation changes in mutant hESCs compared with wt hESCs by Infinium MethylationEPIC BeadChip assay. **c,** Genomic location of differentially methylated sites (DMS), for CpG sites methylated in wt and mutant hESCs. Odds ratio and 95% confidence intervals are displayed in wt and mutant hESCs, comparing their localization in different genomic locations as provided by Illumina (TSS1500, TSS200, 5′UTR, 1stExon, Body, Exon boundaries [ExonBnd], and 3′UTR). Odds ratios were computed against the general distribution of the 684,838 CpGs of our dataset. Indicated 282 out of 684,838 CpG sites are differentially methylated in seven mutant hESCs compared with wt hESCs, 246 out of 286 are associated certain gene. **d,** The distribution of DMRs between wt hESCs and mutant hESCs. 321 genomic regions are differentially methylated (P < 0.05) (in total 4,905 regions from UCSC refgene). **e,** Enrichment of DMRs according to genic regions (left) and distance from CpG island (right) (with a CpG shore, CpG shelve and open sea located 1-2 kbp, 2-4 kbp and >4 kbp away from the nearest CpG island, respectively) between wt and mutant hESCs. 401 genomic regions are differentially methylated (P < 0.05) (in total 4,905 regions). **f, g,** Gene Ontology (GO) categories of genes differentially methylated between wt and mutant hESCs (top categories ranked according to p-value). GO enrichment analysis by DAVID using DMPs associated 246 genes, and DMRs associated 125 genes.

**Extended Data Fig. 6.**
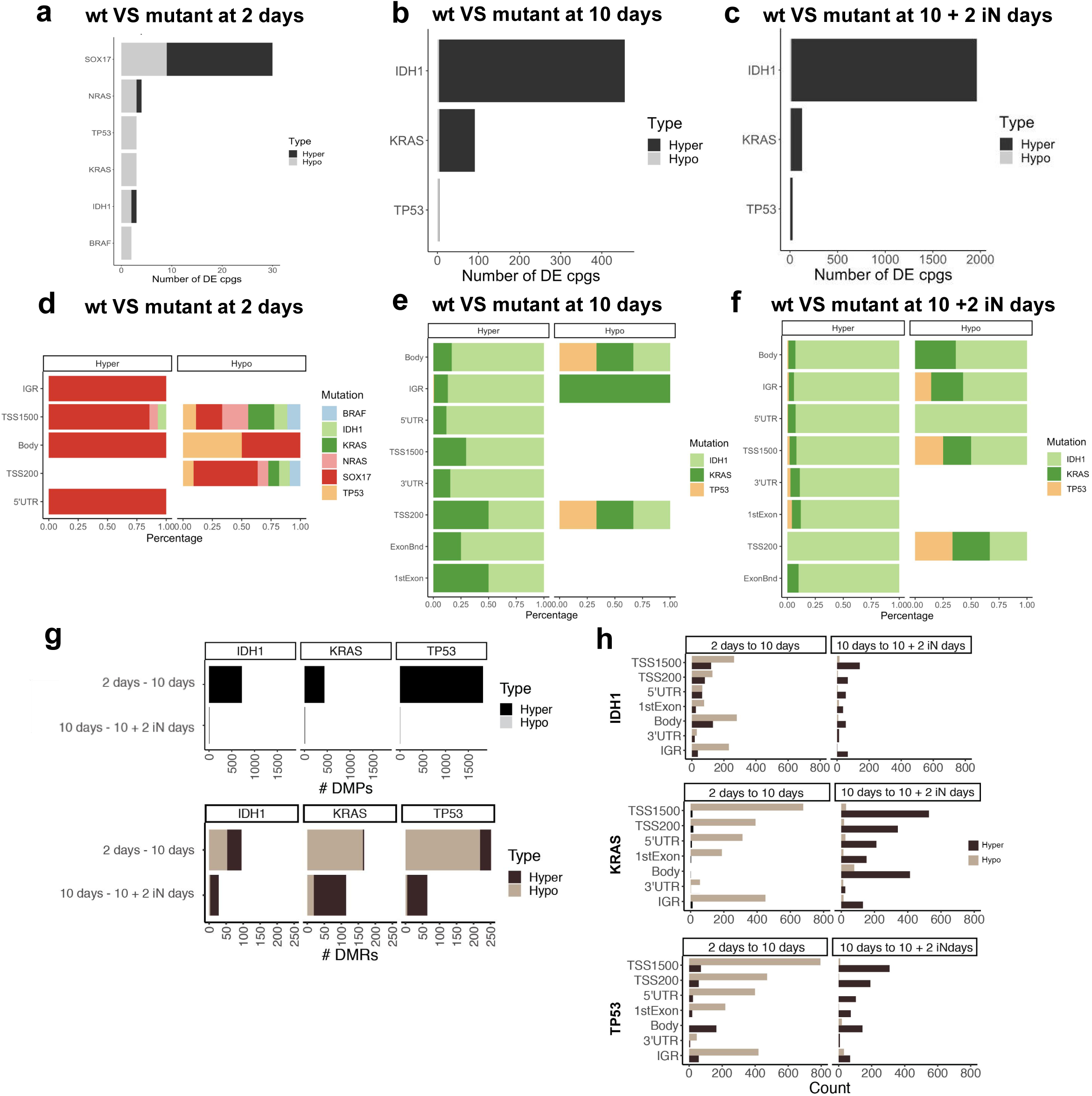
Oncogenic mutations are associated with epigenetic alteration in NPCs and iN cells. **a, b, c,** Number of hypermethylated (beta-value > 0.8 and logFC > 0.3) and hypomethylated (beta-value < 0.2, logFC < -0.3) DMPs in mutant cells compared to wt cells at 2 days, 10 days, and 10 days plus 2 days of neural induction. **d, e, f,** Bar plot shows the percentage of genomic location each DMP was found in mutant cells compared to wt cells at 2 days, 10 days, and 10 days plus 2 days of neural induction (p-value < 0.01). **g,** Number of DMPs and DMRs at 2 to 10 days and 10 to 10 + 2 iN days comparisons for mutant cells (IDH1, KRAS, and TP53). DMPs found in the same comparison for wt cells were removed from the total DMP count. **h,** Number of DMS within the DMRs in (G) across the IDH1, KRAS, and TP53 mutation cells. Black or grey color bar plots represent DMRs found in hyper- or hypomethylated regions.

**Extended Data Fig. 7.**
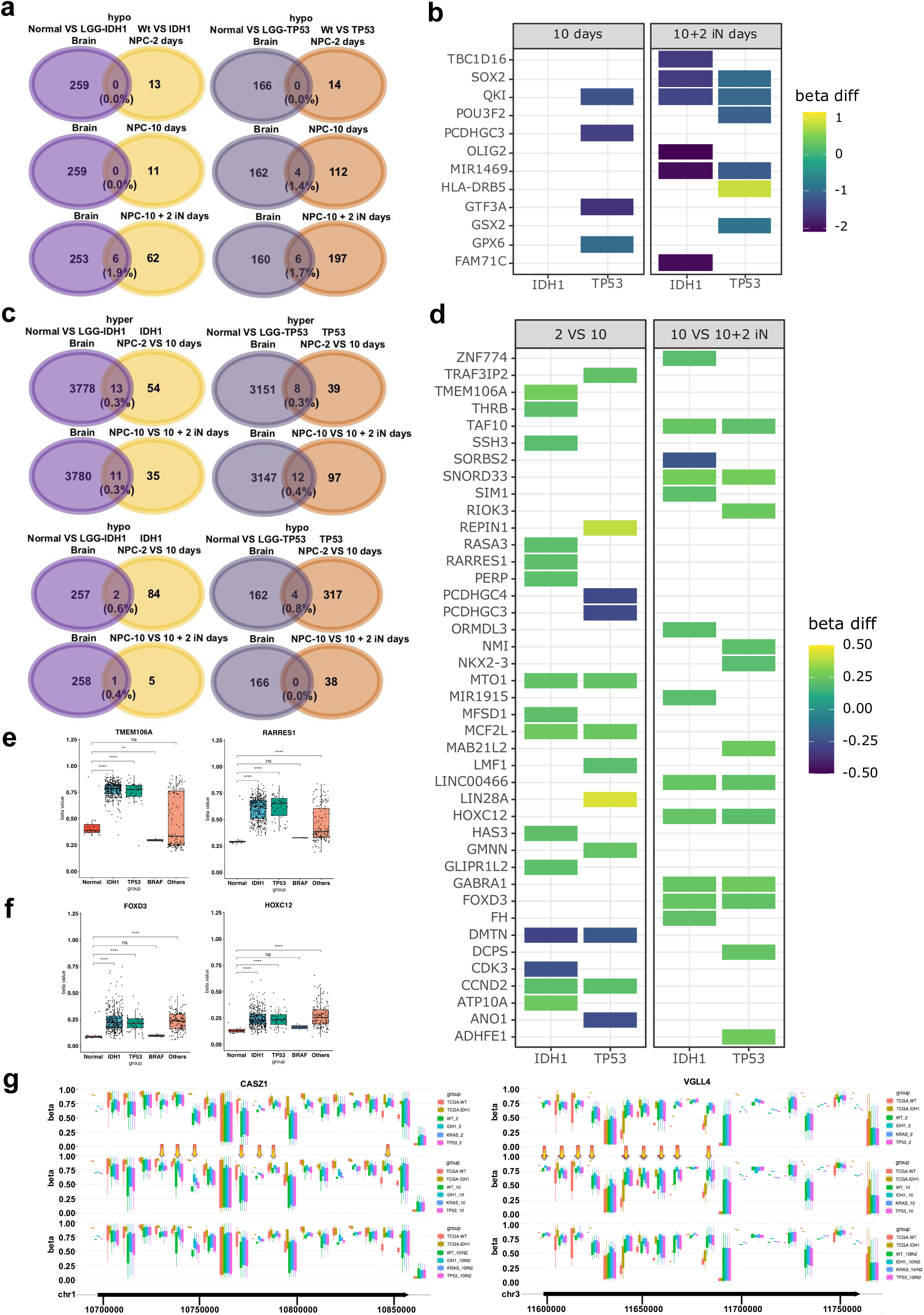
Stem cells with oncogenic mutations share similar epigenetic patterns with brain tumors. **a,** Venn diagram representing the number of hypomethylated genes found in DMRs that come from the control brain VS low-grade gliomas (LGGs) with IDH1 or TP53 mutation, and wt cells VS IDH1 or TP53 mutation cells at 2, 10, and 10 + 2 iN days (p-value < 0.01). **b,** Heatmap of differentially hypomethylated genes found within the overlap node from extended data Fig. 7a at 2, 10, and 10 + 2 iN days (p-value < 0.01). The color scale indicates methylation level from high (yellow) to low (blue). **c,** Venn diagram representing the number of genes found in DMRs that come from the control brain VS LGGs with IDH1 (TP53) mutation, and wt cells VS IDH1 (TP53) mutation cells, at 2 to 10 days comparisons and 10 to 10 + 2 iN days comparisons (p-value < 0.01). **d,** Heatmap of differentially methylated genes found within the overlap node from extended data Fig. 7c at 2 to 10 days comparisons and 10 to 10 + 2 iN days comparisons (p-value < 0.01). The color scale indicates methylation level from high (yellow) to low (blue). **e, f,** From comparing the epigenetic states in LGGs with IDH1-R132H, TP53-R273C, BRAF-V600E, other mutations and control brain cells as measured by DNA methylation from the TCGA dataset, the box plot demonstrates the distribution of DNA methylation of hypermethylated genes in cells at 2 to 10 days comparisons and 10 to 10 + 2 iN days comparisons. (NS: p > 0.05, *: p <= 0.05, **: p <= 0.01, ***: p <= 0.001, ****: p <= 0.0001). **g,** Distribution of DNA methylation marks at CASZ1 and VGLL4 gene loci. The arrows indicate differentially methylation sites.

